# Decoupled neoantigen cross-presentation in tumors with high intratumor heterogeneity reduces dendritic cell activation to limit anti-tumor immunity

**DOI:** 10.1101/2022.12.16.520773

**Authors:** Kim B. Nguyen, Christopher J. Copeland, Coralie M. Backlund, Nory G. Klop-Packel, Tanaka Remba, Byungji Kim, Nishant K. Singh, Michael E. Birnbaum, Darrell J. Irvine, Stefani Spranger

## Abstract

Cancer immunotherapies, in particular checkpoint blockade immunotherapy (CBT), can induce control of cancer growth, with a fraction of patients experiencing durable responses. However, the majority of patients currently do not respond to CBT and the molecular determinants of resistance have not been fully elucidated. Mounting clinical evidence suggests that the clonal status of neoantigens (NeoAg) impacts the anti-tumor T cell response. High intratumor heterogeneity (ITH), where the majority of NeoAgs are expressed subclonally, is correlated with poor clinical response to CBT and poor infiltration with tumor-reactive T cells. However, the mechanism by which ITH blunts tumor-reactive T cells is unclear. We developed a transplantable murine lung cancer model to characterize the immune response against a defined set of NeoAgs expressed either clonally or subclonally to model low or high ITH, respectively. Here we show that clonal expression of a weakly immunogenic NeoAg with a relatively strong NeoAg increased the immunogenicity of tumors with low but not high ITH. Mechanistically we determined that clonal NeoAg expression allowed cross-presenting dendritic cells to acquire and present both NeoAgs. Dual NeoAg presentation by dendritic cells was associated with a more mature DC phenotype and a higher stimulatory capacity. These data suggest that clonal NeoAg expression can induce more potent anti-tumor responses due to more stimulatory dendritic cell: T cell interactions. Therapeutic vaccination targeting subclonally expressed NeoAgs could be used to boost anti-tumor T cell responses.

## Introduction

Engaging tumor-reactive immune responses has been an incredibly powerful tool in the fight against cancer.^1, 2^. Cytotoxic CD8^+^ T cells recognize peptides on class I major histocompatibility complexes (MHCI) expressed on tumor cells, and following recognition mediate specific lysis of their target cell ^3, 4^. While CD8^+^ T cells can recognize many tumor-associated antigens, peptides specific to tumor cells are best suited to drive the most powerful anti-tumor responses ^5, 6^. Amongst the tumor-specific antigens, the class of so-called neoantigens (NeoAg) is best understood thus far. NeoAgs are predominantly derived from non-synonymous mutations in highly expressed protein coding transcripts within the tumor cells ^7^. It has been shown that patients responding to checkpoint blockade immunotherapy (CBT) often experience an expansion in NeoAg reactive T cells within tumor-infiltrating T cells as well as in circulation ^8, 9^. Further, adoptive cell transfer of NeoAg-specific T cells is highly beneficial ^10, 11^ and vaccination can induce objective responses towards tumor-specific NeoAgs ^12–15^.

The presence of CD8^+^ T cells within the tumor microenvironment (TME) is widely accepted as a positive prognostic marker of response to CBT and overall survival ^16^. Over the past years many studies have aimed to establish a correlation between the presence of NeoAgs and CD8^+^ T cells within the tumor. Despite enormous efforts, several independent reports suggest that NeoAg load alone cannot predict response to CBT ^17, 18^. Of note, it seems the prognostic value of NeoAg burden depends on the baseline presence of a T cell infiltrate ^17, 18^. This can be best illustrated in cancer types with high mutational burden, such as melanoma, non-small cell lung cancer, and colon cancer. In those cancer types a sizable proportion of patients lack a productive T cell infiltrate, despite an abundance in predicted NeoAgs ^17–19^. Past studies have indicated that alterations in tumor cell-intrinsic signaling pathways can mediate poor T cell infiltration, typically by means of poor T cell activation or poor T cell recruitment into the TME ^20, 21^. However, these alterations do not account for all patients failing to respond to CBT while harboring high numbers of predicted NeoAgs. Recent studies suggest that intratumor heterogeneity (ITH), which might be highest in patients with high mutational burden, directly impacts the responsiveness to CBT ^22^. Clinical data suggest that clonal NeoAg expression is associated with response to anti-PD-1 CBT in a cohort of patients with NSCLC and with significantly increased overall survival in melanoma patients, following treatment with anti-CTLA-4 antibodies ^22^. In contrast, subclonal NeoAg expression in tumors with high ITH was found to be associated with poor CBT responses and poor CD8^+^ T cell infiltration. These observations were confirmed in a transplantable mouse model using subclones derived from a UVB-irradiated murine melanoma cell line ^23^.

While these initial studies clearly demonstrate that high ITH impairs the anti-tumor immune response, the mechanisms of how the anti-tumor immune response is impaired are still unknown. To interrogate the effect of ITH on anti-tumor T cell responses, we generated a syngeneic transplantable murine lung tumor model that enables us to precisely modulate the degree of ITH using naturally-developed NeoAgs. Using two NeoAgs with different degrees of immunogenicity, we elucidated that responses against the weaker NeoAg were potentiated only in the clonal setting. This synergistic effect was established during T cell activation by cross-presenting conventional type I dendritic cells (cDC1), which acquired a more mature phenotype if they presented both antigens. Intriguingly, RNA-based vaccines targeting the weak NeoAg augmented immune responses in tumors with high ITH, highlighting the potential therapeutic value of targeting weakly immunogenic subclonal NeoAgs.

## Methods

### Mice

C57BL/6 were purchased from Taconic Biosciences and Jackson Laboratory. Rag2^-/-^ and Batf3^-/-^ mice were purchased from Jackson Laboratory and bred in-house. B6 CD45.1 were purchased from Jackson Laboratory. All mice were housed and bred under specific pathogen free (SPF) conditions at the Koch Institute for Integrative Cancer Research Building animal facility. For all studies mice were gendered-matched and age-matched to 6-12 weeks old at the start of experiments. All experimental animal procedures were approved by the Committee on Animal Care (CAC/IACUC) at MIT.

### Tumor cell lines and tissue culture

KP1233 and RMA-S were gifts from the Jacks lab at MIT. KP6S was subcloned from KP1233. Tumor cell lines were cultured at 37°C and 5% CO_2_ in culture media [DMEM (Gibco) supplemented with 10% heat-inactivated FBS (Atlanta Biologicals), 1% penicillin/streptomycin (Gibco), and 20 mM HEPES (Gibco)].

### Functional lentiviral titration

KP6S were plated in a 6-well plate to have 20-40%confluency the following day. Lentivirus was serially diluted and cells were transduced as described here. 48 hours later cells were selected with puromycin or blasticidin (Gibco) and continued selection for 6-8 days until colonies were visible in the wells with the highest dilution. Supernatant was removed and cells were washed once with PBS. 0.5 mL of 10% formalin (Sigma-Aldrich) was added to the wells to fix cells at room temperature for 5 min. Cells were washed with PBS. 0.5 mL of crystal violet stain (0.05% w/v) was added to stain the cells at room temperature for 30 minutes. Cells were washed twice with RODI water. Plates were drained and dried overnight. Blue colonies were counted to determine functional titers.

### Generation of NeoAg-expressing cell lines

KP6S were transduced with the pLV-EF1a-mCherry-mcs lentiviral constructs expressing different NeoAg(s) at an MOI of 0.1. Lentivirus diluted in culture media and supplemented with 4 μg/mL protamine sulfate (Sigma Aldrich, stock is 1 mg/mL in PBS) was added to cells. The media was changed the following day. Selection began 48 hours after cells were transduced. Flow cytometry was used to confirm and quantify construct expression.

### Predicted binding affinities

Peptide sequences were entered into NetMHC Server 4.0 (https://services.healthtech.dtu.dk/service.php?NetMHC-4.0) to determine binding affinities.

### Peptide synthesis

All peptides were synthesized by Genscript at >95% purity with unmodified N- and C-termini.

### RMA-S MHCI stabilization assay

RMA-S cells were cultured in RPMI (Gibco) supplemented with 10% heat-inactivated FBS (Atlanta Biologicals), 1% penicillin/streptomycin (Gibco), and 20 mM HEPES (Gibco)]with 55 μM 2-Mercaptoethanol (Gibco). Cells were placed in a tissue culture incubator and incubated at 28°C the day before the experiment. Cells were collected and counted and resuspend in suspension cell media at 10×10^6^ cells/mL. 100 μL of cells were added to a 96-well plate. A titration of peptide was added to the cells to bring the volume to 200 μL. Cells were incubated for 2 hours at 28°C before antibody staining for flow cytometry analysis.

### Short peptide vaccination

A single dose consisted of 10 μg of peptide (peptide stock is 10 mg/mL resuspended in DMSO) was added to 25 μg cyclic-di-GMP (stock is 1 mg/mL resuspended in PBS) (Invivogen tlrl-nacdg) and PBS added to a final volume of 50 μL. Mice were briefly anesthetized (isoflurane) and the vaccine was administered s.c. at the base of the tail. Mice were given a second identical boost 10 days later and spleens were collected 11 days after the boost.

### Tumor outgrowth studies

Tumor cells were collected by trypsinization (Gibco) and washed 3 times with 1X PBS (Gibco). Cells were resuspended in PBS, and 1×10^6^ tumor cells were injected subcutaneously into the flanks of mice. Subcutaneous tumor area measurements (calculated as length x width) were collected 2-3 times a week using digital calipers until the endpoint of the study.

### DNA extraction and qRT-PCR

DNA was extracted using the Sigma-Aldrich GenElute Mammalian Genomic DNA Miniprep kit following manufacturers’ instructions. Extracted DNA was quantified by NanoDrop. DNA was diluted to yield stock concentrations of 50-120 ng/μL. This was further diluted 1:100 or 1:1000 for the reactions. For each plate, a standard was plated using genomic DNA extracted from cell lines along with genomic DNA extracted from tumor tissue. 20 μL reactions were ran [10 μL 2X SYBR Green PCR Master Mix (Applied Biosystems), 200 μM forward primer, 200 μM reverse primer, 6 μL diluted DNA]. Reactions were run on the StepOne Real-Time PCR System (Applied Biosystems) and CT values were used to determine the amount of DNA contributed by a clone in a single sample. The single cell suspension implanted into mice on day 0 (Input) was used as a normalization factor:

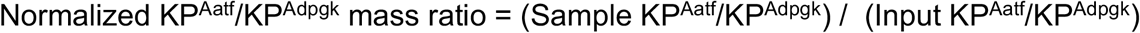

### Tumor dissociation

Tumors were dissected from mice, weighed, and collected in RPMI (Gibco) containing 250 μg/mL Liberase (Sigma-Aldrich) and 1 mg/mL DNase (Sigma-Aldrich). Tumors were minced with dissection scissors or a razor blade and incubated for 45 min at 37°C for enzymatic digestion. Following the digestion, tumor pieces were mashed through a 70 μm filter with a 1 mL syringe plunger to generate a single cell suspension. The dissociated cells were washed 3 times with chilled PBS containing 1% heat-inactivated FBS and 2 mM EDTA (Gibco).

### Flow cytometry

Prior to staining, cells were washed with FACS staining buffer (chilled PBS containing 1% FBS and 2 mM EDTA). If cells were used for intracellular staining Brefeldin A at 1X (BioLegend) was added to all reagents up until the fixation/permeabilization step. Cells were stained for 15 min on ice with eBioscience Fixable Viability Dye eFluor 780 to distinguish live and dead cells and with anti-CD16/CD32 (clone 93, BioLegend) to prevent non-specific antibody binding. Cells were washed once and cell surface proteins were stained for 30 min on ice with fluorophore-conjugated antibodies. Following surface staining cells were washed twice and analyzed directly or fixed with IC Fixation Buffer (eBioscience) for 20 min at RT for analysis the next day. For intracellular staining, cells were washed twice in wash buffer (eBioscience) and incubated with fluorophore-conjugated antibodies for at least 30 minutes or overnight at 4C. Cells were washed twice with FACS staining buffer before running samples. To obtain absolute counts of cells, Precision Count Beads (BioLegend) were added to samples following manufacturer’s instructions. All used antibodies are listed in Suppl. Table 1. Flow cytometry sample acquisition was performed on BD LSRFortessa cytometer and BD Symphony cytometer, and the collected data was analyzed using FlowJo v10.5.3 software (TreeStar).

### Mouse IFN*γ*-ELISpot

All ELISpot specific reagents are part of the IFN*γ*-ELISpot kit from BD Biosciences (Cat 551083). ELISpot plates were coated overnight at 4°C with anti-IFN*γ* antibody. Plates were washed and blocked with DMEM supplemented with 10% FBS, 1% penicillin/streptomycin, and 20 mM HEPES for 2 hr at room temperature (RT). Spleens were harvested from mice and mashed through a 70 μm filter with a 1 mL syringe plunger to generate a single cell suspension. Red blood cells were lysed with 500 μL of ACK Lysing Buffer (Gibco) on ice for 5 min and splenocytes were washed 3 times with chilled PBS. For IFN*γ*-ELISpot assays using peptide restimulation, 1×10^6^ splenocytes were assayed per well in the presence or absence of 10 μg of peptide. As a positive control, splenocytes were incubated with a mixture of 100 ng/mL PMA (Sigma-Aldrich) and 1 μg/mL ionomycin (Sigma-Aldrich). Following an overnight incubation at 37°C and 5% CO2, plates were developed using the BD mouse IFN*γ*-ELISpot kit, following manufacturer’s protocol.

### Irradiated tumor cell vaccination

Tumor cells were trypsinized, washed once in 1X PBS, passaged through an 18g needle to generate a single cell suspension, and further washed twice in 1X PBS. Cells were resuspended at a concentration of 15×10^6^ cells/mL in 1X PBS. Cells were placed in a conical and irradiated with 40 Gy (gray) on ice. Cells were injected into mice immediately following irradiation. Mice were vaccinated as described in short peptide vaccination with both Aatf and Adpgk peptides included in the dose. Spleens were collected for IFN*γ*-ELISpot on day 21 post implantation of lethally irradiated tumor cells.

### Immunotherapeutic modulation

For checkpoint blockade therapy 100 μg each of anti-CTLA-4 (Clone UC10-4F10-11, Bio X Cell BP0032) and anti-PD-L1 (Clone 10F.9G2, Bio X Cell BP0101) or 100 μg each of isotype controls (Clones N/A and LTF2, Bio X Cell BP0091 and BE0090) was administered i.p. diluted in a total volume of 100 μL or 200 μL of PBS on day 7, 10, 13 and 16 following tumor implantation. For CD40 agonism 100 μg of anti-CD40 (Clone FGK4.5, Bio X Cell BP0016-2) was administered i.p. diluted in a total volume of 100 μL of PBS on day 4 following tumor implantation.

### RNA vaccination

Self-replicating RNA based on Venezuelan Equine Encephalitis virus replicons were cloned encoding two copies of the cell penetrating peptide (CPP) penetratin attached to the previously defined NeoAg Aatf (TCTTTTATGGCTCCAATAGACCATACTACTATGTCAGAT) separated by GGS cleavable sites (CPP-Aatf-CPP-Aatf) under the subgenomic promoter and prepared by in vitro transcription as previously described^24^. Replicons were formulated in lipid nanoparticles by microfluidic nanoprecipitation. The lipids were composed of N1,N3,N5-tris(3-(didodecylamino)propyl)benzene-1,3,5-tricarboxamide (TT3) ^25^, (6Z,9Z,28Z,31Z)-Heptatriaconta-6,9,28,31-tetraen-19-yl 4-(dimethylamino) butanoate (DLin-MC3-DMA; MedChemExpress), 1,2-dioleoyl-sn-glycero-3-phosphoethanolamine (DOPE; Avanti Polar Lipids), Cholesterol (Avanti Polar Lipids), and 1,2-dimyristoyl-*rac*-glycero-3-methoxypolyethylene glycol-2000 (DMG-PEG2k; Avanti Polar Lipids) at a molar ratio of 10:25:20:40:5. RNA (stored in RNAse-free water) was diluted in 10 mM citrate buffer at pH 3.0 (Alfa Aesar). The lipids and RNA were mixed using the NanoAssemblr Ignite instrument (Precision Nanosystems) operated with the following settings: volume ratio-2:1; flow rate-12 ml/min; waste volume-0 mL. The RNA-loaded LNPs were dialyzed against PBS prior to use. Mice were immunized with 1 μg of replicon RNA in LNPs i.m. in the gastrocnemius muscle.

### Statistical analysis

All statistical analyses were performed using GraphPad Prism (GraphPad).

## Results

### Cancer cells expressing NeoAgs elicit diverse anti-tumor immune responses

Next-generation genome sequencing combined with MHCI binding prediction algorithms and *in vivo* validation have allowed for the identification of bona fide NeoAgs expressed in murine tumor lines including MC38, B16F10, and TRAMP-C1 ^26–28^. Based on their reported immunogenicity, we selected candidate NeoAgs derived from mutated Adpgk, Aatf, and Cpne1 and immunized C57BL/6 mice with short peptides (8mer and 9mer) containing the mutations to validate their immunogenicity. Immunization with the mutant Adpgk peptide induced appreciable expansion of NeoAg-specific T cells while immunization with Cpne1 and Aatf peptides resulted in low or non-detectable T cell responses, respectively (Fig. 1A; Table 1).

**Figure 1.**
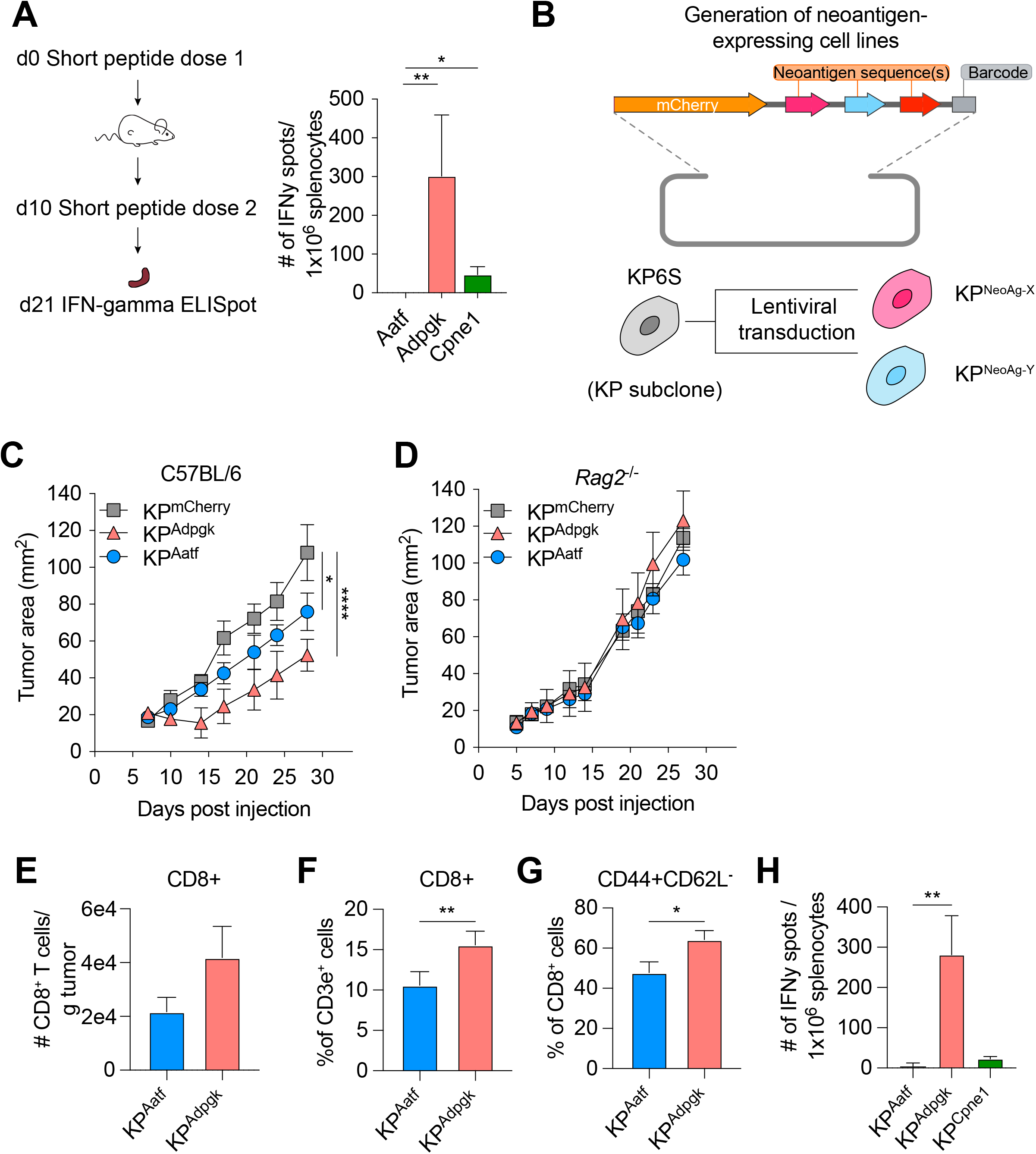
KP6S cell line engineered to express natural NeoAgs elicit variable anti-tumor immune responses. A) Mice were vaccinated with short peptides with cyclic-di-GMP as adjuvant. 10 μg of peptide was delivered s.c. at the base of the tail along with 25 μg of cyclic-di-GMP. An identical dose was delivered s.c. 10 days following the first dose and spleens were collected at day 21 for IFN*γ* ELISpot. Quantification of IFN*γ* producing cells after restimulation from two independent experiments shown as mean ± s.e.m (n=3 per group per experiment). B) Schematic of the lentiviral construct used to transduce the KP6S subclone. C-D) Mice were injected subcutaneously with 1×10^6^ tumor cells in B) WT mice or C) *Rag2^-/-^* mice. Representative data from one of two individual experiments are shown (n=3 or 4 per group per experiment). Quantification of E) absolute numbers of CD8+ TIL per gram tumor from six independent experiments (pooled n=20 per group), F) proportion of CD8+ TIL at day 9 or 10 after tumor-implantation from seven independent experiments (pooled n=23 per group), G) proportion of CD44+CD62L-T_effector_ from seven independent experiments (pooled n=23 per group), H) IFN*γ* producing cells restimulated 9 or 10 days after tumor-implantation using ELISpot from two independent experiments (pooled n=5 per group). *P<0.05, **P<0.01, ****P<0.0001; one-way ANOVA (Kruskal-Wallis) test in A), two-way ANOVA (Tukey) in C-D), Mann-Whitney U in E-H). Data are shown as mean ± s.e.m.

**Table 1.**
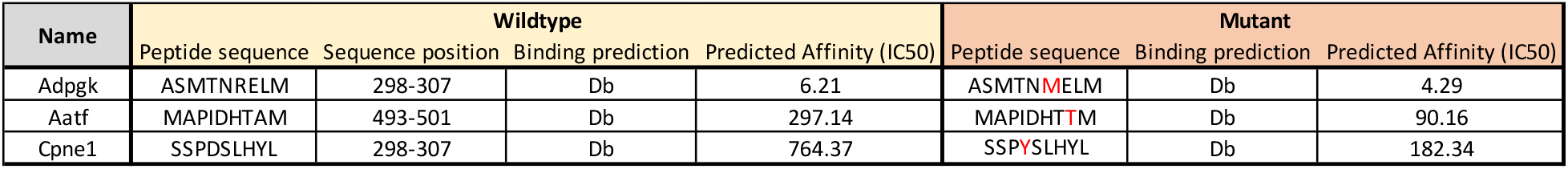
Amino acid sequences of wildtype and NeoAg.

We next generated cell lines to assay anti-NeoAg responses *in vivo* by using KP1233, a lung adenocarcinoma line derived from a Kras^G12D/+^p53^-/-^ mouse ^29^. Because of the inherent cellular heterogeneity observed in many murine cell lines ^30^ we derived a subclone that grew similarly to the parental line in wildtype mice. To drive expression of specific NeoAgs, we expressed one or two NeoAgs linked at the C-terminus of the fluorescent protein mCherry followed by a barcode (Fig. 1B). Subcutaneous implantation of the subclones expressing a single NeoAg (KP^Adpgk^, KP^Aatf^, KP^Cpne1^) indicated that KP^Adpgk^ exhibited early tumor control before growing out while KP^Aatf^ and KP^Cpne1^ cell lines grew out progressively (Fig. 1C, Suppl. Fig. 1A). To confirm that the initial control observed against KP^Adpgk^ tumor cells was mediated by an adaptive immune response, we implanted all cell lines into *Rag2^-/-^* mice and observed that all three cell lines grew progressively with similar kinetics (Fig. 1D, Suppl. Fig. 1B).

Naturally arising NeoAgs expressed in human cancer encompass both highly immunogenic and poorly immunogenic sequences. Thus, we chose the Adpgk, Aatf, and Cpne1 NeoAgs to capture the diversity of NeoAg-specific responses observed in humans ^31^. Analysis of tumor-infiltrating T cells in KP^Adpgk^ and KP^Aatf^ showed that KP^Adpgk^ tumors had a greater degree of infiltration with CD8^+^ T cells compared to KP^Aatf^ tumors (Fig. 1E-F). Further, CD8^+^ T cells in KP^Adpgk^ tumors were more activated based on CD44 staining (Fig. 1G). Additionally, IFN*γ* ELISpot showed a greater peripheral expansion of NeoAg-specific T cells in mice implanted with KP^Adpgk^ tumors compared to KP^Aatf^ or KP^Cpne1^ tumors (Fig. 1H). Assessing the T cell infiltrate in KP^Cpne1^ and KP^mCherry^ tumors revealed that Cpne1-expressing tumor cells were not highly immunogenic as neither CD8+ T cell infiltration nor activation were significantly different between both tumors (Suppl. Fig. 1C-E). The immune responses observed corresponded with MHCI binding affinities predicted by NetMHC 4.0 ^32, 33^ with the mutant Adpgk peptide predicted to have an IC50 of 4.29 nM while the other mutant peptides had IC_50_ values ranging from 90.16 nM (Aatf) to 182.34 nM (Cpne1) (Suppl. Fig. 1F). MHCl-stabilization assays also provided evidence that predicted binding affinities captured the range of peptide-MHCI (pMHCI) affinities for our selection of NeoAgs (Suppl. Fig. 1G). Thus, we established a model of transplantable syngeneic murine tumor lines that express NeoAgs with varying degrees of immunogenicity.

### Homogenous expression of NeoAgs increases the immunogenicity of cancer cells

To assess the impact of different NeoAg expression patterns in tumors we first generated a cell line that expressed both Adpgk and Aatf, hereafter termed KP-Het^Low^ (Fig. 2A). To model heterogeneous NeoAg expression patterns (KP-Het^High^), we inoculated a mixture of 50% KP^Aatf^ cells and 50% KP^Adpgk^ cells (Fig. 2A). We implanted 1×10^6^ cells of KP-Het^High^ and KP-Het^Low^ tumors into mice and observed drastically increased control of tumor outgrowth of KP-Het^Low^ tumors compared to single antigen expressing tumors. In contrast KP-Het^High^ grew progressively, with similar kinetics as observed for KP^Aatf^ (Fig. 2B). In fact, quantitative PCR analysis of KP-Het^High^ tumors showed progressive outgrowth of the Aatf subclone that completely dominates the tumor by day 14 post-implantation (Suppl. Fig. 2A). The control of KP-Het^Low^ tumors was completely lost in *Rag2^-/-^* mice (Fig. 2B), indicating that tumor control was mediated by an adaptive immune response. We also found that the change in tumor cell composition of KP-Het^High^ tumors at later timepoints observed in wildtype mice was also absent in *Rag2^-/-^* mutants. Instead KP^Aatf^ and KP^Adpgk^ cells were maintained at nearly a 1:1 ratio (Suppl. Fig. 2B), providing evidence for immunoediting in this model. These data indicate that adaptive immune responses are capable of controlling tumors with homogenous NeoAg expression while immune editing enables escape of non-immunogenic clones in tumors with heterogeneous NeoAg expression.

**Figure 2.**
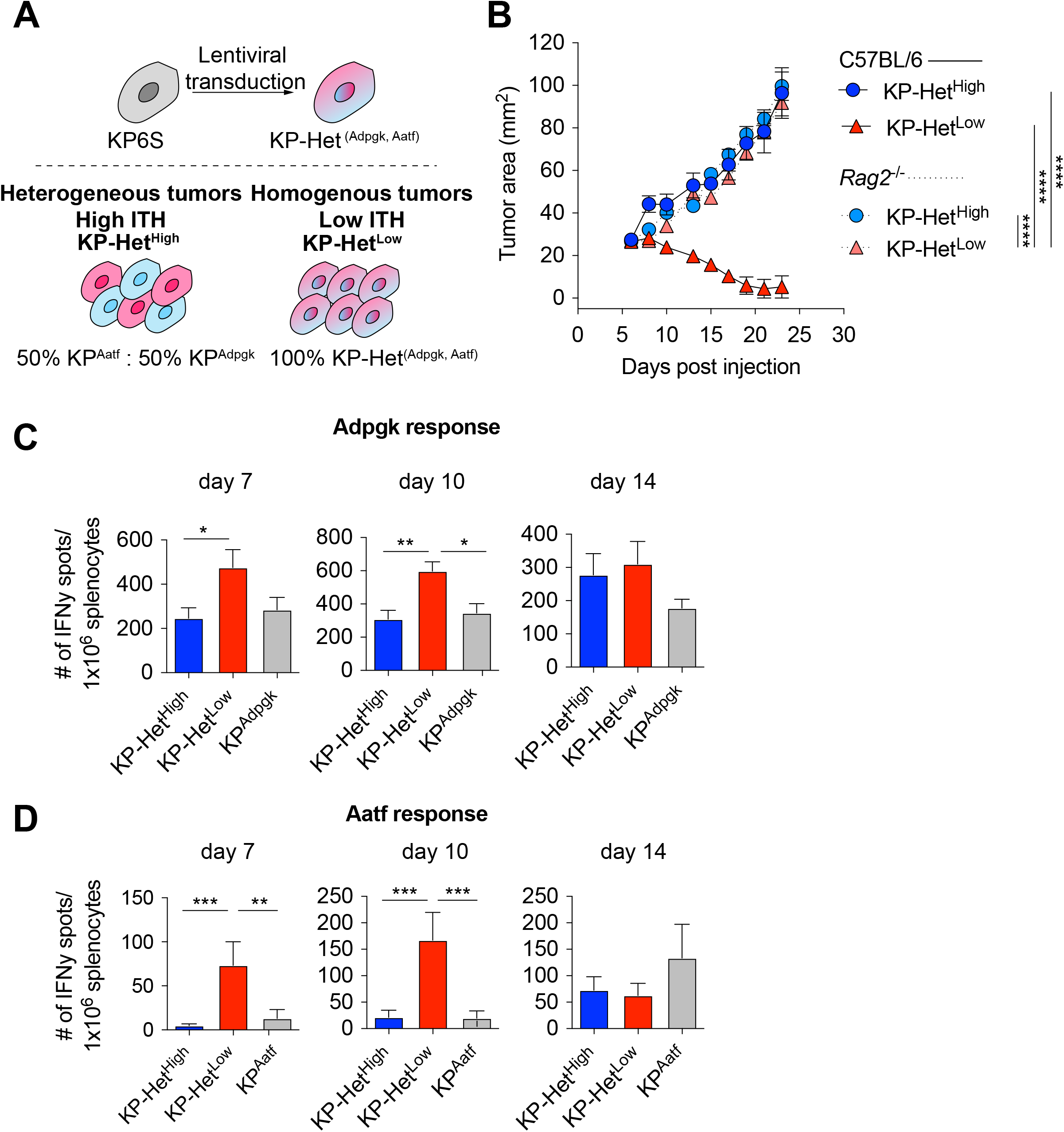
Tumors expressing a pair of NeoAgs homogenously have increased immunogenicity. A) Schematic of the generation of tumors used in B). B) Tumor growth of KP-Het^High^ and KP-Het^Low^ in WT and *Rag2^-/-^* mice. Representative data from one of two individual experiments are shown (n=3 per group per experiment). C-D) Splenocytes from tumorbearing mice were used in an IFN*γ* ELISpot to determine the frequency of NeoAg-specific T cells in the periphery at day 7, 10 and 14 after tumor implantation. Quantification of the C) Adpgk-specific response and D) Aatf-specific response. Pooled data from five independent experiments for day 7 and 10 (n=3 per group per experiment) and three independent experiments for day 14 (n=3 per group per experiment) in A-B). *P<0.05, **P<0.01, ***P<0.001, ****P<0.0001; two-way ANOVA (Tukey) in B), one-way ANOVA (Kruskal Wallis followed by Dunn’s multiple comparisons test) in C-D). Data are shown as mean ± s.e.m.

To obtain insights into the kinetics of the tumor-reactive T cell response, we assayed NeoAg-specific T cells via IFN*γ* ELISpot on days 7, 10 and 14 post tumor implantation. The T cell response towards Adpgk was significantly greater in KP-Het^Low^ tumors compared to KP-Het^High^ tumors at day 7 and greater than both KP-Het^High^ and KP^Adpgk^ tumors at day 10 (Fig. 2C). Strikingly, at day 7 and day 10, the T cell response against Aatf was only detectable in mice implanted with KP-Het^Low^ tumors and was absent in mice bearing KP-Het^High^ or KP^Aatf^ tumors (Fig. 2D). At day 14 the Aatf and Adpgk responses were similar in all tested conditions, suggesting mixed effects of tumor size, antigen availability and loss of functional capacity of T cells over time (Fig. 2B-D). The observed enhanced T cell response against a weakly immunogenic NeoAg is also observed when Cpne1 was co-expressed with Adpgk (Suppl. Fig. 3A-B). We considered the possibility that increasing NeoAg load in a cell could increase immunogenicity by expressing the two weakly immunogenic NeoAgs, Aatf and Cpne1, together. However, this provided no benefit to the Aatf response (Suppl. Fig. 4). These data suggest that homogenous NeoAg expression patterns can increase the peripheral response against poorly immunogenic NeoAgs if they are paired in tandem with a stronger antigen.

### Batf3^+^ dendritic cells are required for anti-tumor responses in KP-Het^Low^ tumors

Given our observation that peripheral T cell responses against weak NeoAgs are enhanced early following tumor inoculation, we postulated that T cell activation of Aatf-reactive T cells in the lymph node might be different between mice bearing KP-Het^Low^ and KP-Het^High^ tumors. While it is established that cross-presenting cDC1 driven by the Batf3 transcription factor are critical for priming CD8^+^-specific responses ^34, 35^, recent work by us and others have also implicated additional cDC subsets ^36^ or compensatory development of Batf3-independent cDC1 ^37^ in mediating anti-tumor immunity. We thus aimed to determine whether Batf3-dependent cDC1 were required for the increased immune control observed against KP-Het^Low^ tumors. We implanted KP-Het^Low^ tumor cells in wildtype, *Rag2^-/-^* and *Batf3^-/-^* mice and observed total loss of tumor control in *Rag2^-/-^* and *Batf3^-/-^* mice (Fig. 3A) indicating that cDC1 are required for the induction of protective T cell responses.

**Figure 3.**
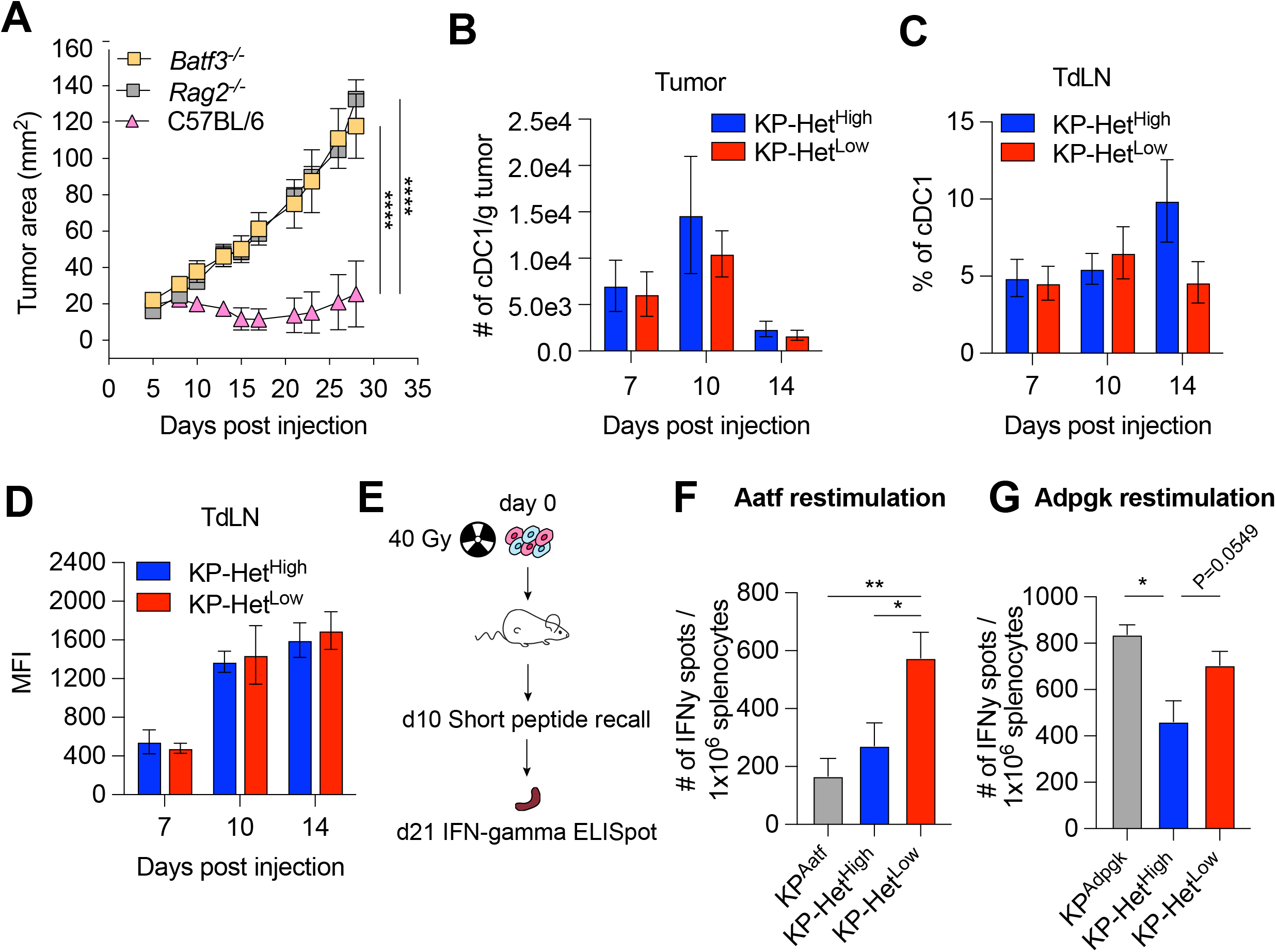
Cross-presenting dendritic cells mediate the increased immunogenicity of KP-Het^Low^ tumors. A) Tumor growth of KP-Het^High^ tumor cells were implanted s.c. into *Batf3^-/-^, Rag2^-/-^* and WT mice. Representative data from three independent experiments (n=5 per group per experiment). B) Number of cDC1 in KP-Het^High^ and KP-Het^Low^ tumors on day 7, 10 and 14 after s.c. implantation. Pooled data from two independent experiments is shown (n=3 per group per experiment). C) Proportion of mCherry^+^ cDC1 in tumor draining lymph nodes. Pooled data from two independent experiments is shown (n=3 per group per experiment). D) Median fluorescence intensity of the mCherry signal of cells from B). Pooled data from two independent experiments for day 7 and 10 and three independent experiments for day 14 is shown (n=3 per group per experiment). E) Experimental schematic for F-G). Tumor cells were irradiated with 40 Gy and 1.5×10^6^ total irradiated cells were immediately s.c. injected into mice. A short peptide boost with both peptides and c-di-GMP as adjuvant was given 10 days after and administered s.c. at the base of the day. 21 days after the irradiated cell implantation spleens were collected for ELISpot. F) Peripheral Aatf-specific response. G) Peripheral Adpgk-specific response. Pooled data from three independent experiments are shown (n=3 or 6 per group per experiment) for B-C). *P<0.05, **P<0.01, ****P<0.0001; two-way ANOVA (Tukey) in A), Mann-Whitney U for each time point between the two tumors was assessed in B-D), one-way ANOVA (Kruskal Wallis followed by Dunn’s multiple comparisons test) in F-G). Data are shown as mean ± s.e.m.

cDC1 can impact anti-tumor T cell responses during T cell activation in the tumor draining lymph node (TdLN) or by facilitating recruitment to the tumor ^35^. Since we observed differences in CD8^+^ T cell infiltration between KP-Het^Low^ and KP-Het^High^, we first assessed the number of tumor-infiltrating cDC1. However, while we observed dynamic changes in the absolute numbers of cDC1 over time, no significant difference was found between the two tumor conditions (Fig. 3B, Suppl. Fig. 5). To track cDC1 carrying tumor cell debris we controlled for mCherry expression in all cell lines by assessing the fluorescent intensity using flow cytometry to ensure equal antigen and fluorophore expression (Suppl. Fig. 6). Assessing the number of tumor-cell-debris carrying mCherry+ cDC1 in the TdLN further affirmed that the differences in T cell activation were not driven by a lack migratory cDC1 bringing antigen to the TdLN, as similar frequencies were detected between the two tumor conditions (Fig. 3C, Suppl. Fig. 5). Analysis of the mCherry MFI amongst the mCherry+ cDC1 similarly showed no significant difference between the KP-Het^Low^ and KP-Het^High^ conditions (Fig. 3D), suggesting that neither cDC1 recruitment to the tumor, trafficking to the TdLN nor amount of available antigen can explain the observed differences in T cell activation.

In the homogeneous KP-Het^Low^ setting, it is conceivable that epitope spreading in response to a rapid and strong Adpgk-specific T cell response might lead to an increase in available Aatf antigen. This increase in antigen could explain an increase in activation of Aatf-reactive T cells compared to the KP-Het^High^ setting. To test whether antigen availability alone might explain the differences in T cell response, we inoculated mice with lethally irradiated tumor cells using single-antigen expressing tumor cell lines, or the KP-Het^Low^ and KP-Het^High^ conditions (Fig. 3E). To ensure robust responses, we recalled T cell responses with a mix of purified Adpgk and Aatf short peptides combined with cyclic-di-GMP as an adjuvant 10 days after the initial injection of irradiated tumor cells (Fig. 3E). 11 days post recall, T cell responses were assessed using an IFN*γ* ELISpot assay (Fig. 3E). Consistent with our previous observations, we observed that the Aatf-specific T cell response was dependent on the context of the NeoAg expression patterns, with greater expansion of Aatf-specific T cells in response to KP-Het^Low^ tumor debris compared to either KP^Aatf^ or KP-Het^High^ tumor debris (Fig. 3F). In contrast, we did observe that the Adpgk response was sensitive to lower antigen availability (Fig. 3G), consistent with previous reports on the correlation between antigen availability and strength of T cell response ^38, 39^. In sum, we identified that NeoAg expression patterns are critical for priming responses against weak NeoAgs, while the antigen load impacts responses towards strong NeoAgs.

### NeoAg presentation on dendritic cells mirror NeoAg expression patterns in the TME

It has been shown that the same dendritic cell can uptake debris containing MHCII and MHCI restricted epitopes allowing the DC to interact with the CD4^+^ T cell for licensing to then activate a productive CD8^+^ T cell response ^40^. Similarly, reports suggest that interactions between a DC and CD8^+^ T cells can impact the maturation state of the DC ^41, 42^. We thus considered the possibility that a strong MHCI epitope might act as a “licensing” response to a weaker MHCI epitope when presented on the same DC. To test this notion, we regenerated the KP-Het^Low^ cell lines by expressing Adpgk linked at the C-terminus of ZsGreen (ZsG) while Aatf maintained its expression with mCherry establishing KP-Het^Low(ZsG-Adpgk,Aatf)^, and a corresponding KP^ZsG-Adpgk^ as control (Suppl. Fig. 7A). We confirmed that these tumor cell lines recapitulated the previously observed outgrowth kinetics (Suppl. Fig. 7B). Given that we established the importance of cDC1 for T cell priming, we focused our analysis on this DC subset and used mCherry and ZsGreen as a readout for tumor cell debris engulfment and antigen presentation (Fig. 4A). We first determined the proportion of single-fluorophore or double-fluorophore positive cDC1 in the TdLN at day 7 post tumor implantation and found that in KP-Het^High^ tumors most of the cDC1 carrying detectable debris were either mCherry^+^ or ZsGreen^+^ (Fig. 4B). In stark contrast, most tumor cell-debris positive cDC1 found in the TdLN draining KP-Het^Low^ tumors were double positive for both mCherry+ and ZsGreen^+^ (Fig. 4B). Within the single positive cDC1 subset in both tumors there was a bias toward ZsGreen^+^ cells (Fig. 4C), which could be attributed to the stability of this fluorescent protein ^43^.

**Figure 4.**
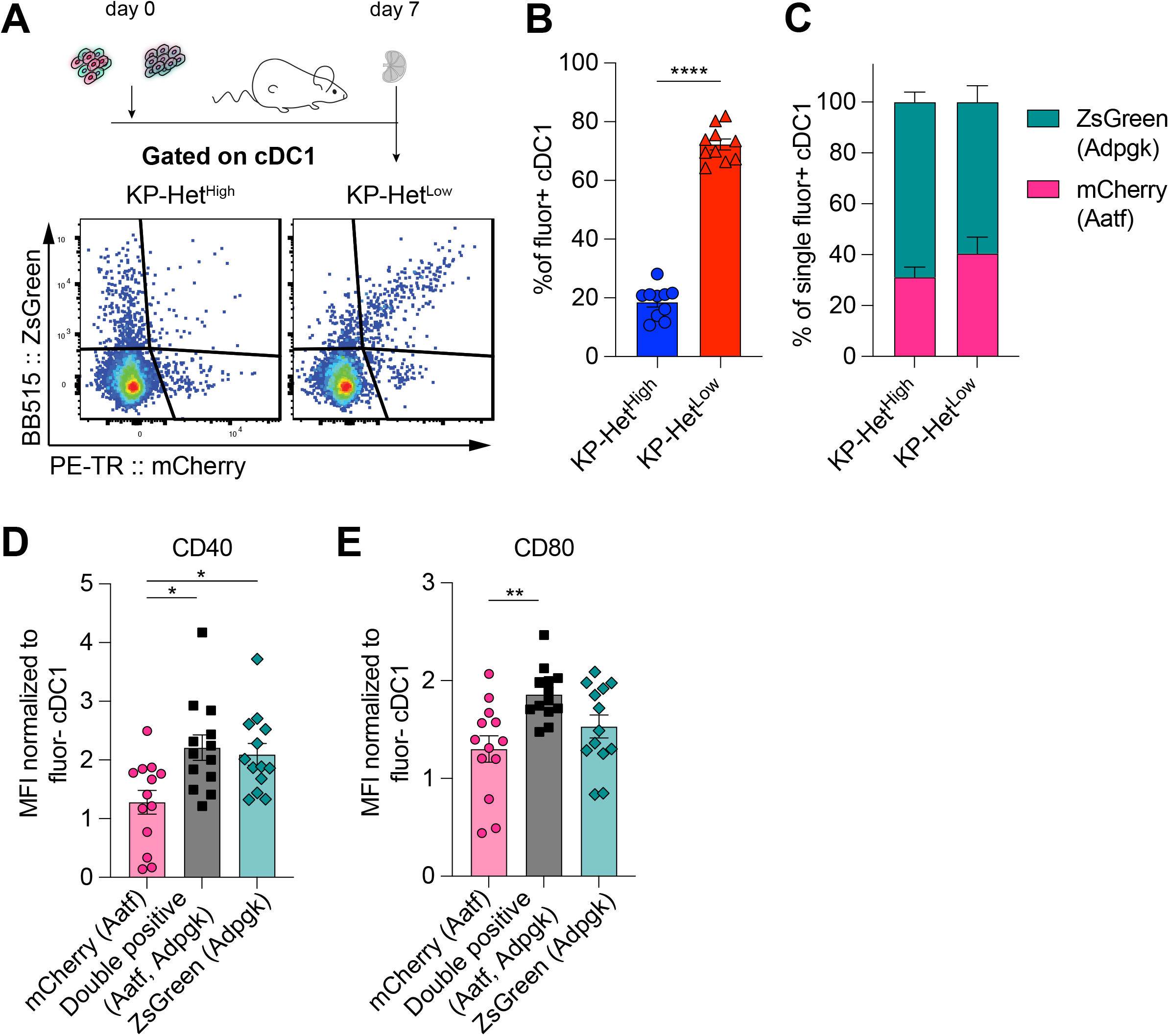
Antigen presentation on dendritic cells in the tumor draining lymph node mirror antigen expression patterns in the TME. A) Experimental schematic for B-C). B) Quantification of the proportion of cDC1 that are double positive (mCherry+ZsGreen+) in tumors. C) Proportion of mCherry+ or Zsgreen+ cDC1 in the single positive population. Pooled data from three independent experiments are shown (n=3 or 4 per group per experiment). D) Normalized CD40 median fluorescence intensity for single positive and double positive populations. E) Normalized CD80 median fluorescence intensity for the same sample populations in D). Pooled data from three independent experiments are shown (pooled n=13 per group) for D) and E). *P<0.05, **P<0.01; one-way ANOVA (Kruskal Wallis followed by Dunn’s multiple comparisons test) in B-E). Data are shown as mean ± s.e.m.

Previous reports have indicated that costimulatory markers were upregulated on dendritic cells following “licensing” interactions with CD4^+^ but also CD8^+^ T cells ^41, 42, 44, 45^. We therefore assessed the expression of CD40 and CD80 on single or double fluorophore positive cDC1 populations in the TdLN. Affirming our initial hypothesis, we observed significantly greater expression of the costimulatory molecules CD40 and CD80 in double-positive cDC1 compared to mCherry^+^ cDC1 that engulf only Aatf debris (Fig. 4D-E). CD40 expression was comparable between Adpgk-ZsGreen^+^ and double positive cDC1, while CD80 was highly expressed on all cDC1 (Fig. 4D-E), suggesting indeed that the Adpgk-specific T cell response might induce upregulation of co-stimulatory molecules. In sum, these findings suggest that the antigen-dependent interaction between cDC1 and Adpgk-specific T cells could result in increased activation (“licensing”) of cDC1, and subsequently an increased capacity of cDC1 to prime Aatf-reactive T cells, if the same cDC1 also presents the weaker NeoAg.

### Prophylactic RNA vaccination expands Aatf-specific T cells and increases response of heterogeneous tumors to CBT

Clinically, a high degree of ITH is associated with poor responses to CBT. To determine whether our established model system faithfully recapitulated resistance to therapy we inoculated KP-Het^Low^ or KP-Het^High^ in C57BL/6 mice and treated mice with dual CBT, consisting of anti-CLTA4 and anti-PD-L1 antibodies on day 7, 10, 13, and 16. Consistent with clinical observations, our model showed that KP-Het^Low^ tumors could be fully controlled following therapy (Fig. 5A). In contrast, KP-Het^High^ tumors, engineered to resemble tumor with high ITH, showed mixed responses (Fig. 5A). Our gained insights into the mechanism of resistance in tumor with heterogeneous NeoAg expression suggest that increased tumor control in homogenous tumor was associated with a more rapid and robust expansion of T cell responses towards weaker NeoAgs. Therefore, we aimed to determine whether prophylactic vaccination might increase tumor control of KP-Het^High^ tumors in the context of dual CBT. Mice were vaccinated with self-amplifying RNA (replicons) encased in a lipid nanoparticle administered intramuscularly (i.m.) and boosted i.m. 2 weeks after the initial dose. Using this strategy, we induced a detectable Aatf-specific response 7 days post boost (Fig. 5B), unlike short peptide vaccinations (Fig. 1A). 1×10^6^ KP-Het^High^ tumor cells were inoculated 7 days post boost and dual CBT was administered on day 7, 10, 13 and 16 post tumor implantation (Fig. 5C). As seen before, KP-Het^High^ tumors showed mixed responses following CBT, ranging from progressive growth to stable disease (Fig. 5D). Prophylactic vaccination alone only resulted in a modest reduction of tumor growth (Fig. 5D). However, we observed a synergistic effect of the combination treatment of prophylactic vaccination and dual CBT, with 2 out of 7 (29%) objective responses with one complete response and one stable disease (Fig. 5D). Thus, we established proof-of-concept that targeting a subclonal, weakly immunogenic NeoAg could be a viable strategy to increase CBT response.

**Figure 5.**
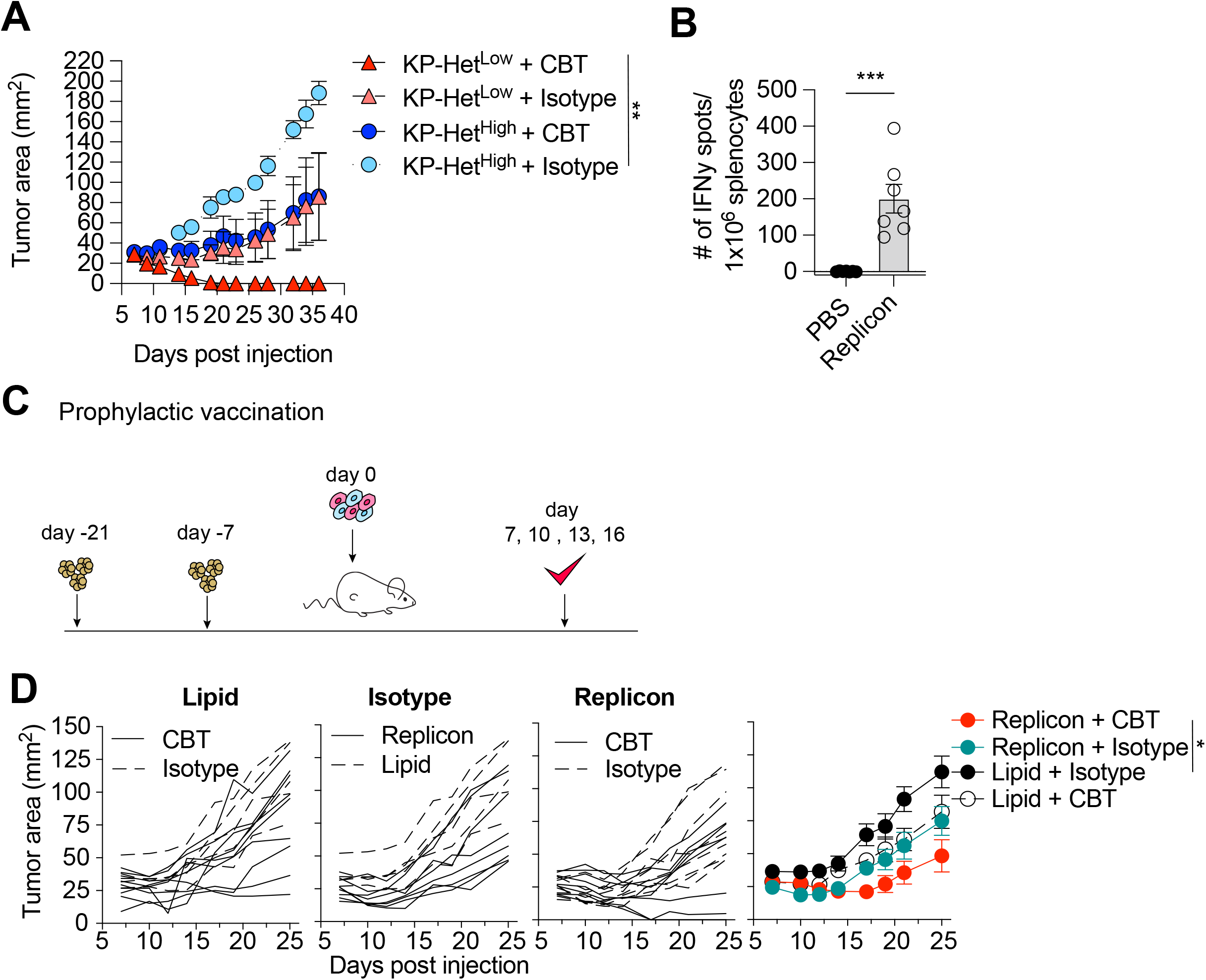
Prophylactic mRNA replicon vaccination increases response to checkpoint blockade therapy in KP-Het^High^ tumors. A) Tumor growth of KP-Het^High^ and KP-Het^Low^ WT mice treated with CBT or control. 100 μg of each antibody (CBT or isotype control) was administered i.p. on day 7, 10, 13, and 16 after implantation. Representative data from one of two individual experiments are shown (n=3 per group per experiment). B) IFN*γ* ELISpot using splenocytes from mice vaccinated with replicons expressing Aatf. C) Experimental schematic for prophylactic vaccination in C). Three weeks before tumor challenge mice are initially vaccinated, replicons are administered intramuscularly (i.m.). Animals are boosted one week before challenge. CBT is administered intraperitoneally (i.p.) on day 7, 10, 13 and 16 following s.c. implantation of KP-Het^High^. D) KP-Het^High^ outgrowth in WT mice. Individual traces for mice dosed with a lipid-only control and treated with CBT or an isotype antibody control, treated only with isotype antibody control and dosed with the replicon or lipid only control, and vaccinated with the replicon and treated with CBT or an isotype control (left to right). Far right plot is the averaged results. Representative data from three independent experiments (n=7 or 10 per group per experiment). *P<0.05, **P<0.01, ***P<0.001; two-way ANOVA (Tukey) in A) and D), Mann-Whitney U in B). Data are shown as mean ± s.e.m.

### Therapeutic RNA vaccination with CD40 agonism synergizes with CBT in KP-Het^High^ tumors

Given that the majority of NeoAgs emerge during tumor development and thus disqualify prophylactic vaccination, we next aimed to determine the utility of therapeutic vaccination with CBT as a therapeutic strategy. Four days after tumor implantation, mice were dosed with RNA replicons and continually dosed every week thereafter (Fig. 6A). Our analysis of the cDC1 compartment in tumors with clonal NeoAg expression further suggested that CD40:CD40 ligand interactions would enhance induction of Aatf-reactive effector T cell responses (Fig. 4D). Furthermore, previous studies have shown that CD40 stimulation could induce strong systemic anti-tumor responses leading to regression of tumors in preclinical models ^46, 47^ and it has also been shown to synergize with CBT ^39, 48^. Thus, we combined the vaccination approach with a single dose of agonistic anti-CD40 antibody along with the first vaccine dose (Fig. 6A). Dual CBT was administered i.p. on day 7, 10, 13 and 16 post tumor implantation (Fig. 6A).

**Figure 6.**
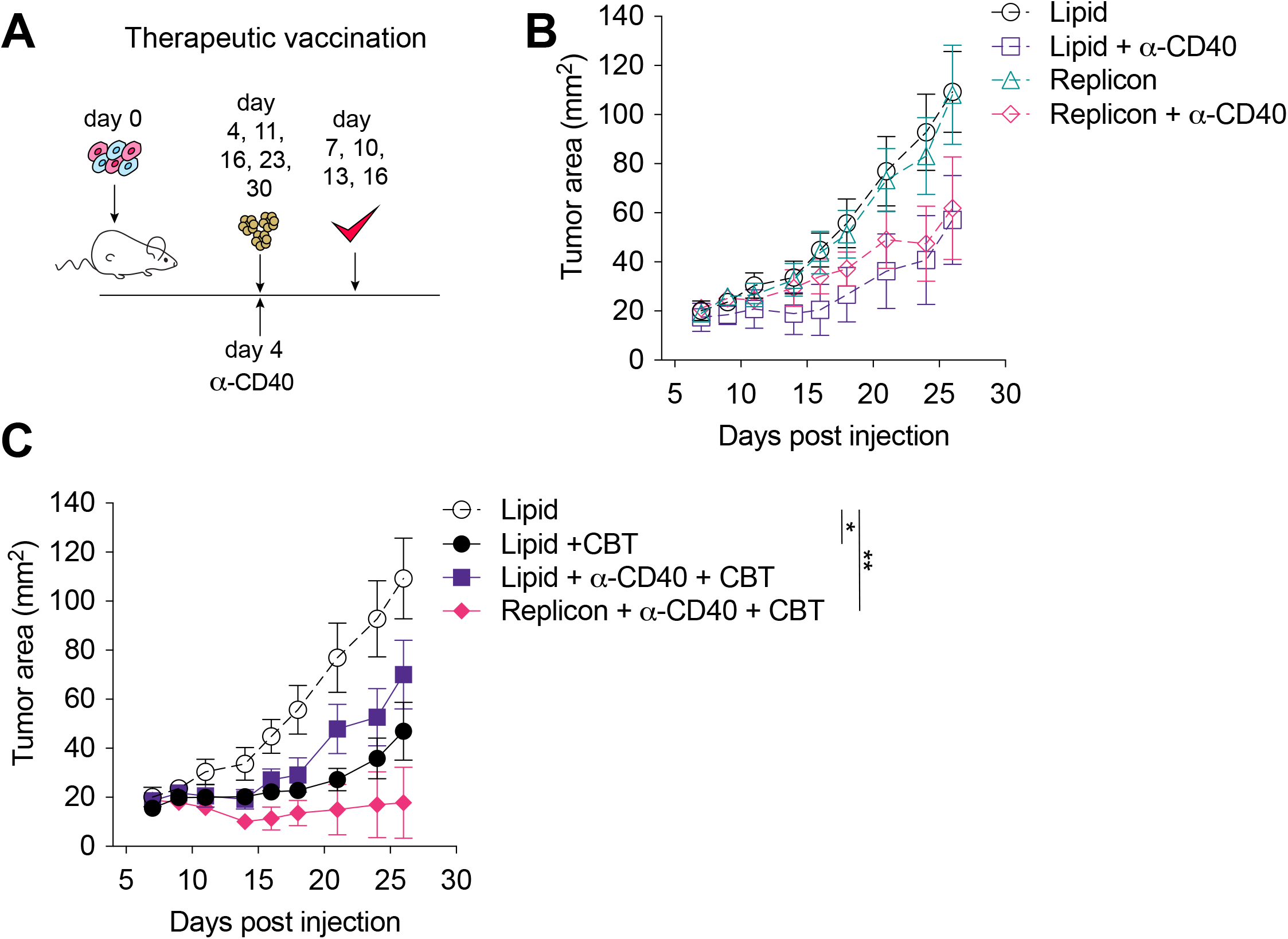
Therapeutic mRNA replicon vaccination synergizes with CBT and CD40 agonism in KP-Het^High^ tumors. A) Experimental schematic for therapeutic vaccination in B and C. Tumor-bearing mice were vaccinated with replicon vaccine (i.m.) on day 4 post tumor inoculation and continually vaccinated every week. Anti-CD40 antibody (100 μg) was given with first vaccination only. Dual CBT was administered i.p. on day 7, 10, 13 and 16 post tumor inoculation. B) Tumor growth of KP-Het^High^ treated with replicons or lipid only with or without anti-CD40 antibody. C) Tumor growth of KP-Het^High^ treated with replicons or lipid only with or without anti-CD40 and CBT. n=5 per group. The dotted line is the same cohort observed in B) of mice administered with only lipid and isotype control antibodies. *P<0.05, **P<0.01; two-way ANOVA (Tukey) in B and C). Data are shown as mean ± s.e.m.

Similar to CBT alone, vaccination alone or agonistic CD-40 antibody administration exhibited only a modest response characterized by slowing of tumor growth (Fig. 6B). In stark contrast the triple combination of CBT with therapeutic vaccination and a single dose of agonistic CD40 antibody induced complete tumor control in 3 of 5 (60%) mice and a significant delay in tumor growth in the remaining 40% of mice (Fig. 6C). CD40 agonism with vaccination was able to induce a modest reduction of tumor growth with 1/5 (20%) mice exhibiting complete tumor regression, while CD40 agonism with CBT exhibited mixed response comparable to CBT alone where stable disease was observed in 1/7 (14%) of mice although all tumors eventually progressed (Fig. 6B-C).These data suggest that for therapeutic vaccination against poorly immunogenic antigens, especially in the heterogenous setting, optimal priming can be achieved when the vaccine induced response is augmented with an initial agonistic CD40 treatment and these T cell responses can be sustained using CBT treatment.

## Discussion

Clinical and pre-clinical studies have established that ITH impairs the anti-tumor immune response ^22, 23, 49^, but the mechanism blunting T cell mediated immunity in tumors with heterogeneous NeoAg expression is still unknown. We utilized a reductionist approach to study the effect of heterogeneous NeoAg expression on the resulting anti-tumor immune response. By comparing NeoAgs expressed in subclonal or clonal settings we discovered that immune responses against poorly immunogenic NeoAgs, such as Aatf, were enhanced when clonally expressed with a strong NeoAg. Mechanistically we identified that clonal expression of NeoAgs allows for simultaneous presentation of peptides on the same cross-presenting cDC1, which showed increased expression of costimulatory molecules such as CD40 in the presence of an immunogenic antigen. Therapeutic vaccination against weak NeoAgs demonstrated synergy with CBT only when the first dose was administered along with anti-CD40 agonist antibody. In sum, our data suggest that subclonally expressed NeoAgs elicit weaker anti-tumor T cell responses against less immunostimulatory peptides, but at the same time suggest that these poorly immunogenic peptides might be ideal candidates for therapeutic vaccination in combination with targeted DC maturation.

We observed that migratory cDC1 carrying both antigens to the TdLN showed higher expression levels of the co-stimulatory ligands CD40 and CD80 compared to cDC1 carrying only the weak NeoAg. One might hypothesize that the difference in tumor debris carried by the cDC1 could result in different interactions with T cells against the stronger NeoAg in the TdLN. Our observation of increased CD40 on cDC1 carrying immunogenic NeoAg are consistent with the concept of DC licensing, where a DC carrying both CD4^+^ and CD8^+^ epitopes are being “licensed” by the helper T cell response ^50^. Recent work demonstrated that cDC1 can be licensed by CD4^+^ T cells, which induce a more mature phenotype via interaction of CD40 with CD40L ^51^. By inducing a more mature phenotype “licensed” cDC1 were found to have a greater ability to prime the CD8^+^ T cell response. Further, it was shown that CD40 expression alone on cDC1 results in a more robust expansion of antigen specific CD8^+^ T cells, providing further evidence that increased expression of CD40 might directly impact priming of tumor-reactive T cells ^52^. In our model, dual-debris carrying cDC1 expressed more CD40 than cDC1 carrying Aatf debris, but not more than cDC1 with Adpgk debris. Because the increase in CD40 expression appears to be restricted to Adpgk-debris carrying cDC1, we conclude that Adpgk-reactive T cells induce upregulation of CD40 on antigen carrying cDC1 and act in a similar manner as CD4^+^ helper T cell responses.

For the selection of potent NeoAgs, much emphasis has been put on the binding affinity of the NeoAg peptide to MHC and the relative binding to its wildtype counterpart ^7, 11, 53^. However, these predictions focusing on high affinity NeoAgs rarely predict more than a handful of actionable antigens. Recent studies however have shown that antigen-dominance can dampen the immunogenicity of the subdominant antigen ^54^ resulting in reduced immune-mediated protection. This study focused on the clonal expression of the model antigens SIINFEKL and SIYRYYGL, both binding to H2-Kb with affinities ^54^. Therapeutic vaccination allowed the sub-dominant response against SIYRYYGL to regain its full functional capacity suggesting that improper priming resulted from competition for the same MHC allele during cross-presentation. In our system, however, we observe that the antigen with the higher affinity (Adpgk; 4.3 nM) did provide “help” over competition for binding of H2-Db with the Aatf antigen with a lower affinity of 90 nM. While we did observe that the Adpgk response was dominant, as previously reported ^55^, we did observe a greater expansion of both NeoAg-specific responses when both antigens were clonally expressed. This observation suggests a therapeutic potential for thus far unused NeoAgs with a lower binding affinity to MHC. It is interesting to note that, Aatf was first identified as a potential NeoAg, however was described to be unlikely immunogenic antigen due to its mutant amino acid location near the carboxy terminus ^26^. Our finding underscores the importance of validation of NeoAg immunogenicity *in vivo.* Furthermore, our study highlights the importance of studying NeoAgs with highly variable immunogenicity as this might preclude detrimental effects of immunodominance between multiple high affinity NeoAgs. This might be especially important clinically as the NeoAg landscape in patients currently does contain peptides with predicted weaker affinity ^31^.

Prophylactic and therapeutic vaccination in our model system demonstrated that CD40 agonism may be sufficient for inducing better anti-tumor responses, but not that it is necessary for all antigens. For optimal translation into clinical practice, it will be critical to determine which antigen specific T cell responses would benefit the most from CD40 agonism. For this, it will be critical to fully elucidate the stimulatory capacity of cDC1 carrying high or low affinity NeoAgs or multiple NeoAgs. Further, it will be critical to determine whether NeoAg expression might mimic our observations made for Aatf. Several reports have shown that low antigen levels in murine cancer models resulted in T cell dysfunction and tumor progression ^39, 56–58^. In a colon carcinoma model with low expression of high affinity antigens, it was shown that CD40 agonism synergized effectively with CBT ^39^, suggesting that low antigen abundance during T cell activation might have similar effects as a low affinity NeoAg. Given that most studies assessing the role of CD40 during CD8^+^ T cell activation presume CD4^+^ T cell help ^50^ it will likewise be critical to decipher the differences between CD4^+^ and CD8^+^ helper responses. Conceivable differences between CD4^+^ and CD8^+^ responses could be the levels of produced cytokines, or the repertoire of co-stimulatory ligands involved. Interestingly, CD40L^+^ CD8^+^ T cells were found to be functionally similar to helper CD4^+^ T cells in one study though the levels of IFN*γ* were lower in CD40L^+^ CD8^+^ T cells compared to CD4^+^ helper T cells in their model ^59^. This is consistent with the observation that IFN*γ* produced by CD8^+^ T cells can induce DC maturation, including upregulation of costimulatory molecules, and expression of IL-12p70 ^41, 60, 61^. The importance of IL-12 for the induction of a productive CD8^+^ T cell responses has been shown in many preclinical mouse models at steady state and following therapeutic interventions ^61–64^. Another consequence of DC licensing is that mature DCs can produce chemokines to facilitate recruitment of naïve CD8^+^ T cells, thereby increasing the chances of finding cognate CD8^+^ T cells to activate ^65^. This notion is highly consistent with our observation that Aatf-reactive T cells are detectable at earlier timepoints when expressed homogenously compared to the heterogeneous NeoAg expression pattern. Future studies employing tools to detect NeoAg-presentation on cDC1 and identification of NeoAg-reactive T cells in situ will be needed to delineate how spatial organization affects the priming response.

While our observations are predominately focused on the induction of a NeoAg T cell response, cDC1 mediated restimulation of effector T cells is also a critical feature in the tumor microenvironment ^66, 67^. However, within the tumor the expression patterns of NeoAg will likely be even more critical, as spatial analysis of the tumor suggest the formation of areas of clonal growth resulting in patches of NeoAg expression ^68–70^. Thus, in tumors with heterogeneous NeoAg expression, cDC1 will consequently only present NeoAg from surrounding tumor cells, thus limiting the stimulatory potential for tumorinfiltrating T cells. It is plausible that this process might accelerate the induction of terminal T cell exhaustion.

In sum, our study provides the first mechanistic insights into how heterogeneous NeoAg expression mediates weaker anti-tumor CD8^+^ T cell responses. With an emphasis on the initial T cell priming response, we were able to elucidate that CD8^+^ T cell responses against abundant, high affinity antigens can enhance CD8^+^ T cell responses against lower affinity NeoAgs by increasing the stimulatory capacity of cDC1. This synergistic effect, however, is limited to homogenous NeoAg expression in tumor cells that result in dual cross-presentation on cDC1. Our work underscores the necessity of improving prediction of functional NeoAg immunogenicity in a patient specific context, considering intratumor heterogeneity, expression level and binding affinity. Understanding these parameters, their dependencies, and their collective impact on the functional immunogenicity of each NeoAg has the potential to enlarge the number of actionable NeoAg for targeted vaccination.

## Acknowledgements

We thank Melissa Duquette for mouse colony maintenance and laboratory support; and Paul Thompson for administrative support. We thank Leon Yim for technical support and Malte Roerden for critical discussions and technical assistance. We thank the Koch Institute Swanson Biotechnology Center for technical support. This work was supported by the Melanoma Research Alliance Young Investigator Award, the A Breath of Hope Lung Foundation, the Koch Institute Frontier Research program, and the Ludwig Center at MIT. K.B.N. is supported by the F31 5F31CA261093-02 (NCI) and S.S. is a Pew-Steward Scholar of the Pew Charitable Trust and holds the Howard S. (1953) and Linda B. Stern Career Development Professorship.

## Author Contributions

K.B.N and S.S. conceptualized the study. K.B.N and S.S. designed experiments and interpreted data. K.B.N, C.C., and N.K.P. performed experiments and analyzed data. N.S. and M.B. generated reagents. C.M.B., T.R., B.J.K., and D.J.I. assisted with vaccination studies. K.B.N. and S.S. wrote and revised the manuscript. S.S. acquired funding. S.S. supervised the study.

---

**Supplemental Figure 1.**
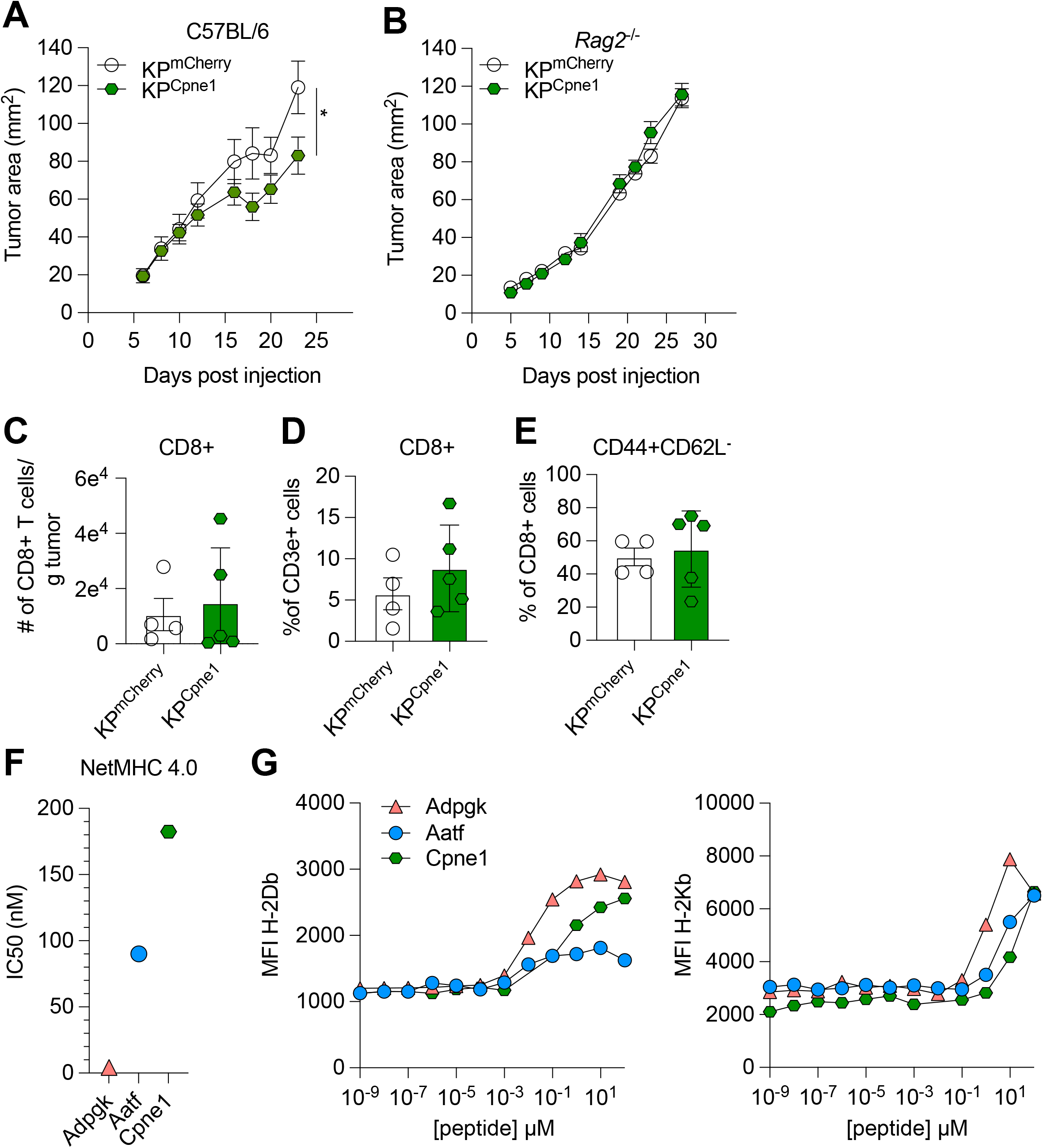
Characterization of the immunogenicity of an array of natural NeoAgs. A-B) Mice were injected subcutaneously with 1×10^6^ tumor cells in A) WT mice or B) *Rag2^-/-^* mice. Representative data from one of two individual experiments are shown (n=5 per group per experiment) in A) and one individual experiment (n=4 per group) in B). Quantification of C) absolute numbers of CD8+ TIL per gram tumor from two independent experiments (pooled n=4 or 5 per group), D) proportion of CD8+ TIL at day 9 or 10 after tumor-implantation from two independent experiments (pooled n=4 or 5 per group), E) proportion of CD44+CD62L-T_effector_ from two independent experiments (pooled n=4 or 5 per group). F) Predicted binding affinity values measured as IC50 values (nM) from NetMHCpan-4.0 server. G) RMA-S cells were incubated with a titration of individual peptides and then stained with antibodies to discern preferential binding to H-2Db (left) or H-2Kb (right). *P<0.05; two-way ANOVA (Tukey) in A-B), Mann-Whitney U in C-E). Data are shown as mean ± s.e.m.

**Supplemental Figure 2.**
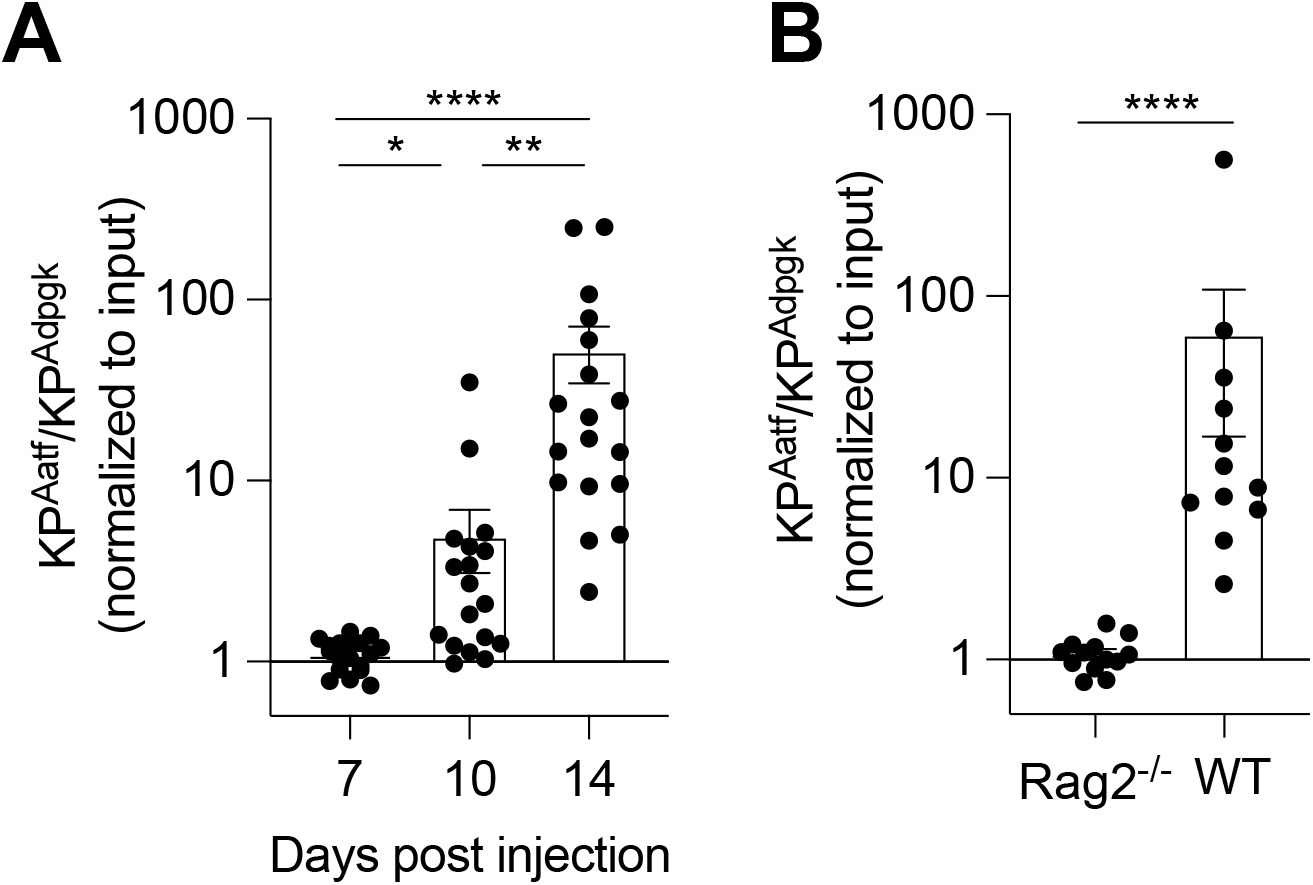
KP-Het^High^ tumors are immune edited. A-B) 1×10^6^ KP-Het^High^ cells were implanted in WT or *Rag2^-/-^* mice. A) Tumors from WT mice were collected at day 7, 10, 14 and 17 post tumor inoculation for genomic DNA extraction used for qRT-PCR to determine tumor composition. Pooled data from three independent experiments (pooled n=18 for each group). B) Tumors implanted in WT or *Rag2^-/-^* mice collected at day 14 or 17 post-tumor implantation. Pooled data from 3 independent experiments (pooled n=12 for WT, pooled n=13 for *Rag2^-/-^).* *P<0.05, **P<0.01, ****P<0.0001; one-way ANOVA (Kruskal-Wallis followed by Dunn’s multiple comparisons) in A), Mann-Whitney U in B).

**Supplemental Figure 3.**
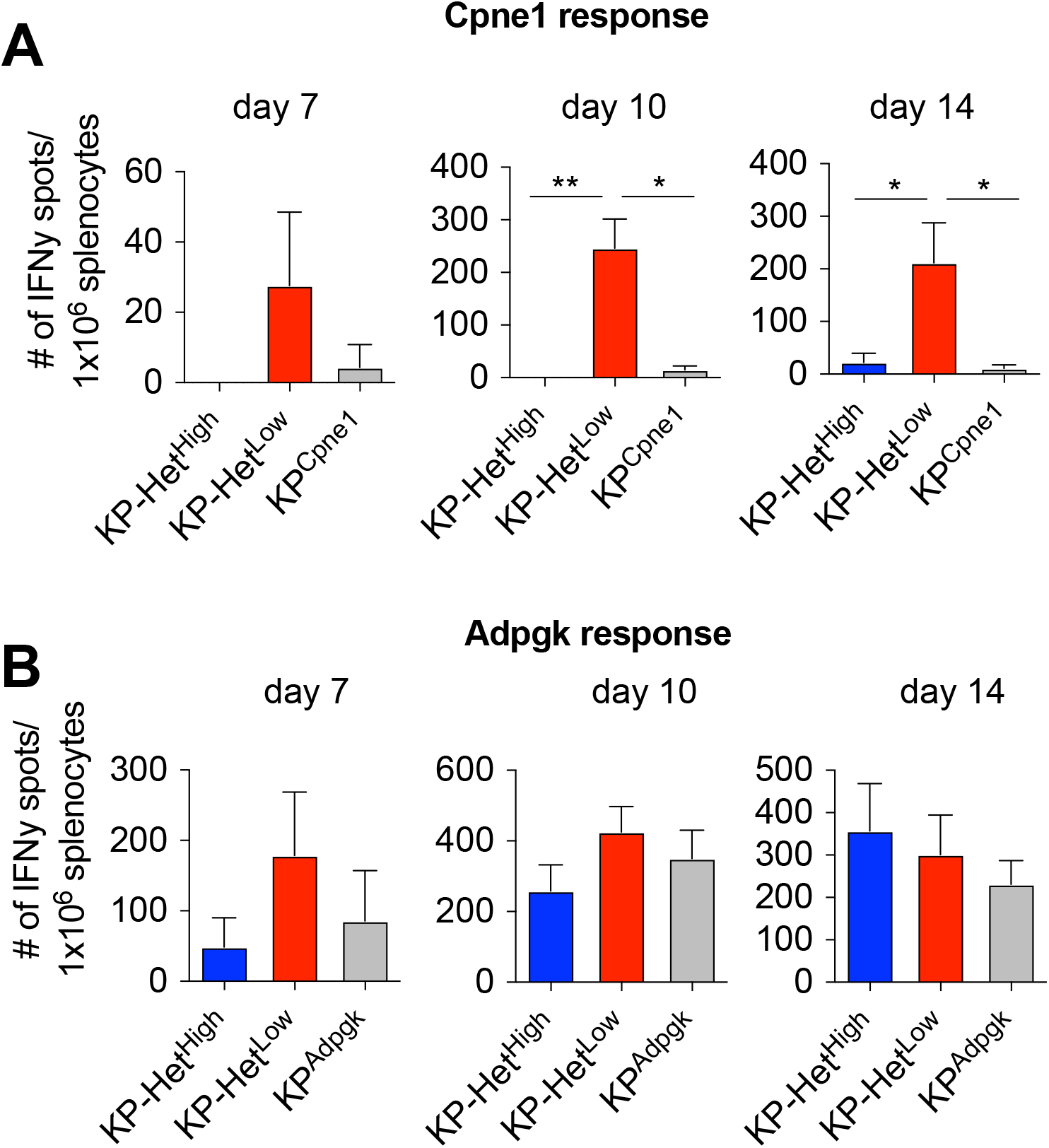
Expansion of NeoAg-specific T cells directed against weak antigens occurs earlier when they are co-expressed with a stronger antigen. Splenocytes from tumor-bearing mice expressing Adpgk and/or Cpnel NeoAgs were used in an IFN*γ* ELISpot to determine the frequency of NeoAg-specific T cells in the periphery at day 7, 10 and 14 after tumor implantation. Quantification of the A) Cpne1-specific response and B) Adpgk-specific response. Pooled data from two independent experiments (n=3 per group per experiment) in A-B). *P<0.05, **P<0.01; one-way ANOVA (Kruskal Wallis followed by Dunn’s multiple comparisons test) in A-B). Data are shown as mean ± s.e.m.

**Supplemental Figure 4.**
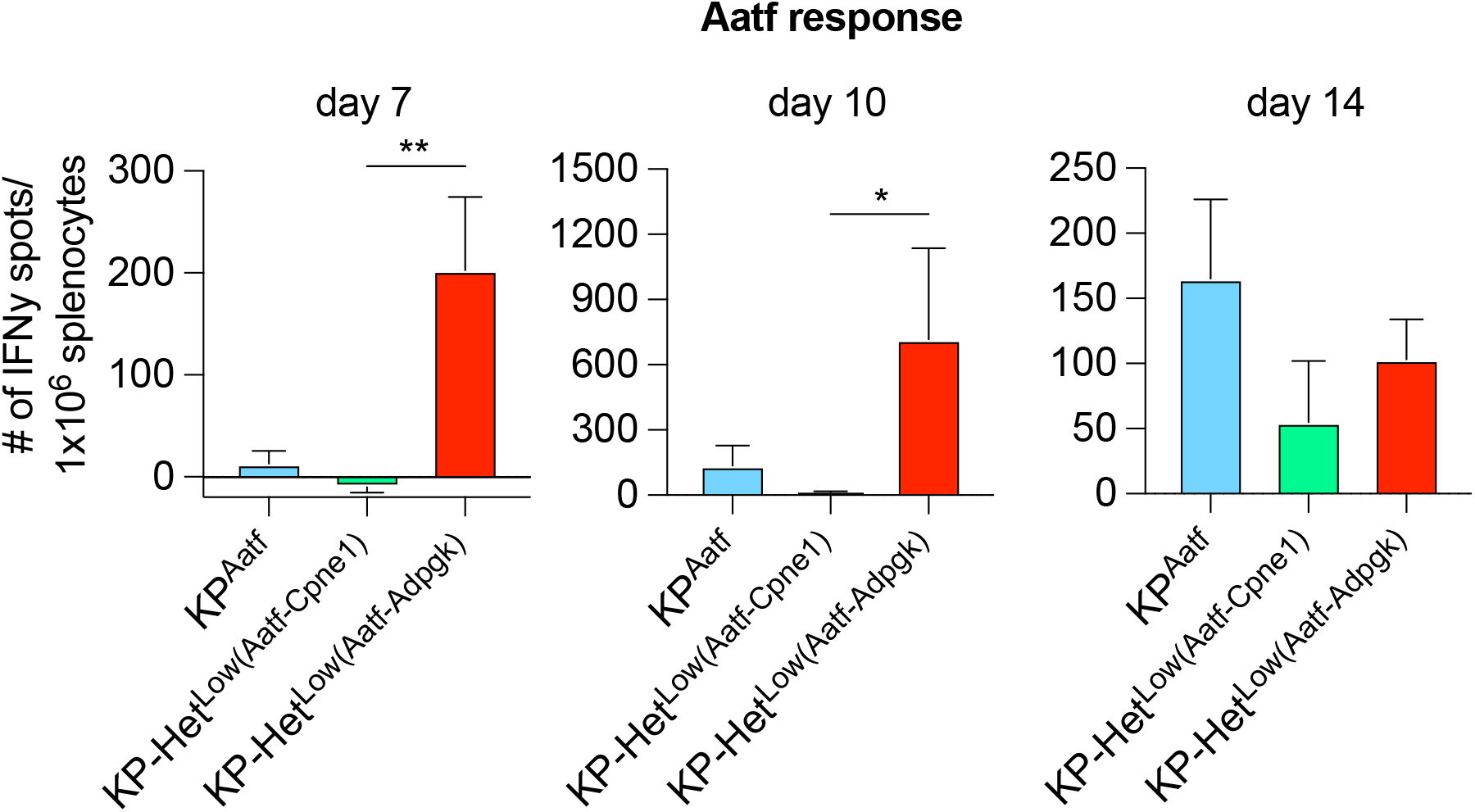
Increased Aatf-specific T cell expansion is not due to increased numbers of antigens expressed in KP-Het^Low^ tumors. Splenocytes from clonal tumor-bearing mice were used in an IFN*γ* ELISpot to determine the frequency of NeoAg-specific T cells in the periphery at day 7, 10 and 14 after tumor implantation. Quantification of the Aatf-specific response. Pooled data from two independent experiments (n=3 per group per experiment). *P<0.05, **P<0.01; one-way ANOVA (Kruskal Wallis followed by Dunn’s multiple comparisons test) in A). Data are shown as mean ± s.e.m.

**Supplemental Figure 5.**
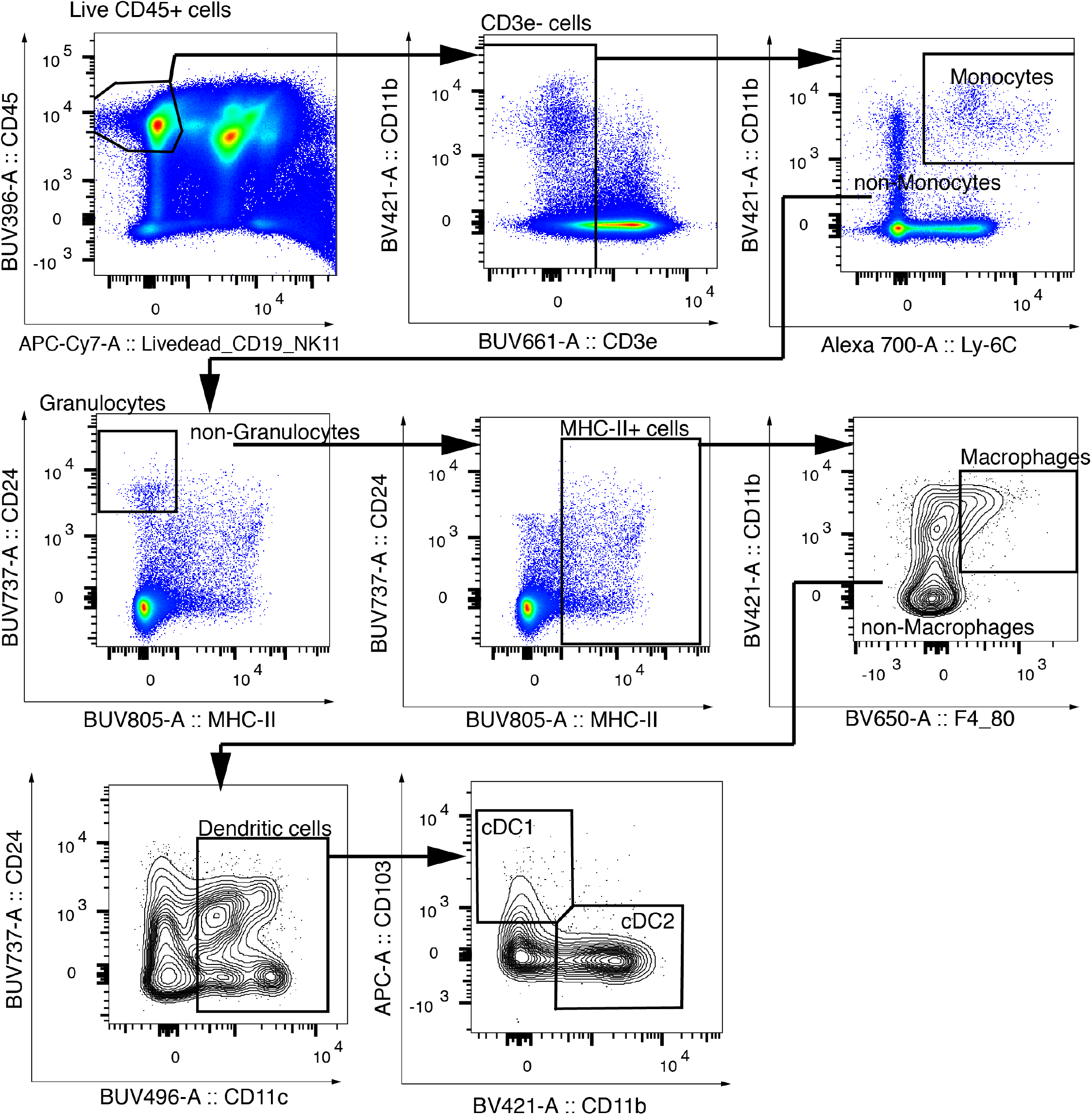
Gating strategy for cDC1. Representative gating strategy to identify cDC1 in the TdLN used in Figure 3 and 4.

**Supplemental Figure 6.**
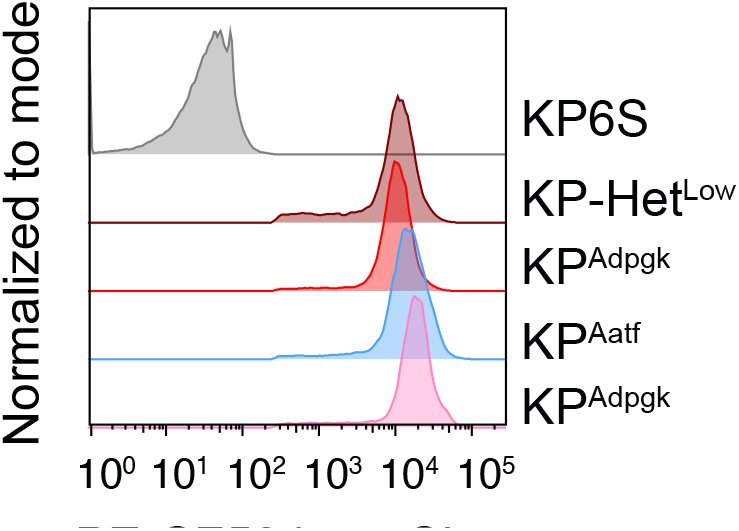
KP cell lines express similar levels of NeoAg. Representative examples of a flow cytometry analysis of mCherry expression in all cell lines used.

**Supplemental Figure 7.**
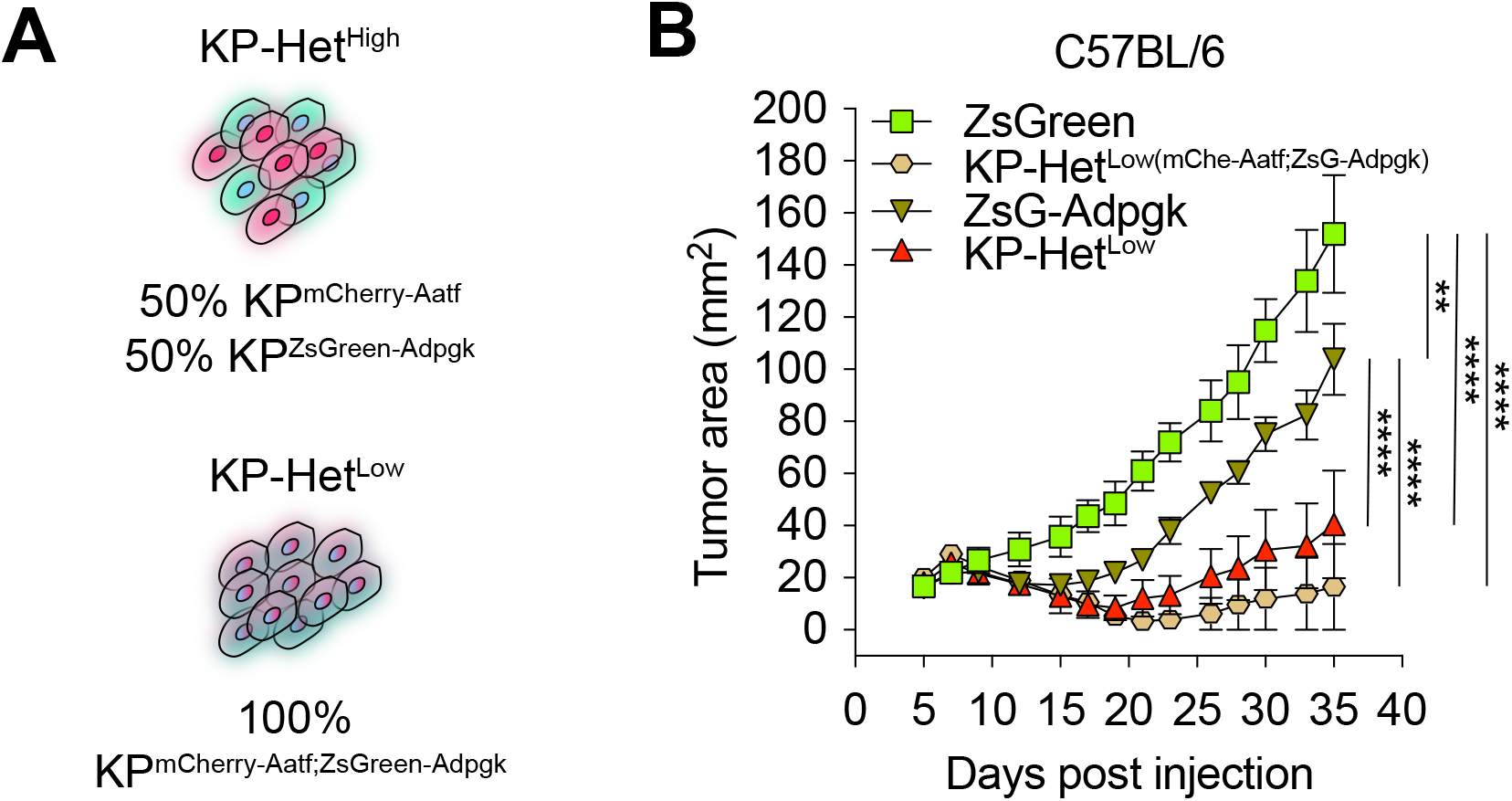
Clonal expression of Adpgk and Aatf NeoAgs results in increased immunogenicity regardless of linked or separate expression of NeoAgs in the same cell. A) Schematic of cells generated used in B). B) Tumor growth in WT mice. n=3 per group. **P<0.01, ****P<0.0001; two-way ANOVA (Tukey) in A). Data are shown as mean ± s.e.m.

## References

1. Waldman AD, Fritz JM, Lenardo MJ. A guide to cancer immunotherapy: from T cell basic science to clinical practice. Nature Reviews Immunology 2020 20:11. 2020;20(11):651–68. doi: 10.1038/s41577-020-0306-5.

2. Esfahani K, Roudaia L, Buhlaiga N, Del Rincon VS, Papneja N, Miller WH. A review of cancer immunotherapy: from the past, to the present, to the future. Current Oncology. 2020;27(Suppl 2):S87. doi: 10.3747/CO.27.5223.

3. Stinchcombe JC, Bossi G, Booth S, Griffiths GM. The immunological synapse of CTL contains a secretory domain and membrane bridges. Immunity. 2001;15(5). doi: 10.1016/S1074-7613(01)00234-5.

4. Isaaz S, Baetz K, Olsen K, Podack E, Griffiths GM. Serial killing by cytotoxic T lymphocytes: T cell receptor triggers degranulation, re-filling of the lytic granules and secretion of lytic proteins via a non-granule pathway. European journal of immunology. 1995;25(4). doi: 10.1002/eji.1830250432.

5. Schietinger A, Philip M, Schreiber H. Specificity in cancer immunotherapy. Seminars in immunology. 2008;20(5). doi: 10.1016/j.smim.2008.07.001.

6. Minati R, Perreault C, Thibault P. A Roadmap Toward the Definition of Actionable Tumor-Specific Antigens. Frontiers in Immunology. 2020;11. doi: 10.3389/fimmu.2020.583287.

7. Schumacher TN, Scheper W, Kvistborg P. Cancer Neoantigens. Annual review of immunology. 2019;37. doi: 10.1146/annurev-immunol-042617-053402.

8. van Rooij N, van Buuren MM, Philips D, Velds A, Toebes M, Heemskerk B, van Dijk LJ, Behjati S, Hilkmann H, El Atmioui D, Nieuwland M, Stratton MR, Kerkhoven RM, Kesmir C, Haanen JB, Kvistborg P, Schumacher TN. Tumor exome analysis reveals neoantigen-specific T-cell reactivity in an ipilimumab-responsive melanoma. Journal of clinical oncology: official journal of the American Society of Clinical Oncology. 2013;31(32):e439–42. doi: 10.1200/JCO.2012.47.7521. PubMed PMID: 24043743; PMCID: PMC3836220.

9. Riaz N, Havel JJ, Makarov V, Desrichard A, Urba WJ, Sims JS, Hodi FS, Martín-Algarra S, Mandal R, Sharfman WH, Bhatia S, Hwu WJ, Gajewski TF, Slingluff CL, Chowell D, Kendall SM, Chang H, Shah R, Kuo F, Morris LGT, Sidhom JW, Schneck JP, Horak CE, Weinhold N, Chan TA. Tumor and Microenvironment Evolution during Immunotherapy with Nivolumab. Cell. 2017;171(4):934–49.e16. doi: 10.1016/J.CELL.2017.09.028.

10. Robbins PF, Lu YC, El-Gamil M, Li YF, Gross C, Gartner J, Lin JC, Teer JK, Cliften P, Tycksen E, Samuels Y, Rosenberg SA. Mining exomic sequencing data to identify mutated antigens recognized by adoptively transferred tumor-reactive T cells. Nature Medicine 2013 19:6. 2013;19(6):747–52. doi: 10.1038/nm.3161.

11. Verdegaal EME, De Miranda NFCC, Visser M, Harryvan T, Van Buuren MM, Andersen RS, Hadrup SR, Van Der Minne CE, Schotte R, Spits H, Haanen JBAG, Kapiteijn EHW, Schumacher TN, Van Der Burg SH. Neoantigen landscape dynamics during human melanoma–T cell interactions. Nature 2016 536:7614. 2016;536(7614):91–5. doi: 10.1038/nature18945.

12. Carreno BM, Magrini V, Becker-Hapak M, Kaabinejadian S, Hundal J, Petti AA, Ly A, Lie WR, Hildebrand WH, Mardis ER, Linette GP. A dendritic cell vaccine increases the breadth and diversity of melanoma neoantigen-specific T cells. Science. 2015;348(6236):803–8. doi: 10.1126/SCIENCE.AAA3828/SUPPL_FILE/CARRENO.SM.PDF.

13. Keskin DB, Anandappa AJ, Sun J, Tirosh I, Mathewson ND, Li S, Oliveira G, Giobbie-Hurder A, Felt K, Gjini E, Shukla SA, Hu Z, Li L, Le PM, Allesøe RL, Richman AR, Kowalczyk MS, Abdelrahman S, Geduldig JE, Charbonneau S, Pelton K, Iorgulescu JB, Elagina L, Zhang W, Olive O, McCluskey C, Olsen LR, Stevens J, Lane WJ, Salazar AM, Daley H, Wen PY, Chiocca EA, Harden M, Lennon NJ, Gabriel S, Getz G, Lander ES, Regev A, Ritz J, Neuberg D, Rodig SJ, Ligon KL, Suvà ML, Wucherpfennig KW, Hacohen N, Fritsch EF, Livak KJ, Ott PA, Wu CJ, Reardon DA. Neoantigen vaccine generates intratumoral T cell responses in phase Ib glioblastoma trial. Nature 2018 565:7738. 2018;565(7738):234–9. doi: 10.1038/s41586-018-0792-9.

14. Johanns TM, Miller CA, Liu CJ, Perrin RJ, Bender D, Kobayashi DK, Campian JL, Chicoine MR, Dacey RG, Huang J, Fritsch EF, Gillanders WE, Artyomov MN, Mardis ER, Schreiber RD, Dunn GP. Detection of neoantigen-specific T cells following a personalized vaccine in a patient with glioblastoma. Oncoimmunology. 2019;8(4). doi: 10.1080/2162402X.2018.1561106.

15. Ott PA, Hu Z, Keskin DB, Shukla SA, Sun J, Bozym DJ, Zhang W, Luoma A, Giobbie-Hurder A, Peter L, Chen C, Olive O, Carter TA, Li S, Lieb DJ, Eisenhaure T, Gjini E, Stevens J, Lane WJ, Javeri I, Nellaiappan K, Salazar AM, Daley H, Seaman M, Buchbinder EI, Yoon CH, Harden M, Lennon N, Gabriel S, Rodig SJ, Barouch DH, Aster JC, Getz G, Wucherpfennig K, Neuberg D, Ritz J, Lander ES, Fritsch EF, Hacohen N, Wu CJ. An immunogenic personal neoantigen vaccine for patients with melanoma. Nature. 2017;547(7662):217–21. doi: 10.1038/nature22991. PubMed PMID: 28678778; PMCID: PMC5577644.

16. van der Leun AM, Thommen DS, Schumacher TN. CD8+ T cell states in human cancer: insights from single-cell analysis. Nature reviews Cancer. 2020;20(4):218. doi: 10.1038/S41568-019-0235-4.

17. Mauriello A, Zeuli R, Cavalluzzo B, Petrizzo A, Tornesello ML, Buonaguro FM, Ceccarelli M, Tagliamonte M, Buonaguro L. High Somatic Mutation and Neoantigen Burden Do Not Correlate with Decreased Progression-Free Survival in HCC Patients not Undergoing Immunotherapy. Cancers. 2019;11(12). doi: 10.3390/CANCERS11121824.

18. McGrail DJ, Pilié PG, Rashid NU, Voorwerk L, Slagter M, Kok M, Jonasch E, Khasraw M, Heimberger AB, Lim B, Ueno NT, Litton JK, Ferrarotto R, Chang JT, Moulder SL, Lin SY. High tumor mutation burden fails to predict immune checkpoint blockade response across all cancer types. Annals of Oncology. 2021;32(5):661–72. doi: 10.1016/J.ANNONC.2021.02.006.

19. Spranger S, Luke JJ, Bao R, Zha Y, Hernandez KM, Li Y, Gajewski AP, Andrade J, Gajewski TF. Density of immunogenic antigens does not explain the presence or absence of the T-cell-inflamed tumor microenvironment in melanoma. Proceedings of the National Academy of Sciences of the United States of America. 2016;113(48):E7759–E68. doi: 10.1073/pnas.1609376113. PubMed PMID: 27837020; PMCID: PMC5137753.

20. Nguyen KB, Spranger S. Modulation of the immune microenvironment by tumor-intrinsic oncogenic signaling. The Journal of cell biology. 2020;219(1). doi: 10.1083/jcb.201908224. PubMed PMID: 31816057; PMCID: PMC7039199.

21. Lawson KA, Sousa CM, Zhang X, Kim E, Akthar R, Caumanns JJ, Yao Y, Mikolajewicz N, Ross C, Brown KR, Zid AA, Fan ZP, Hui S, Krall JA, Simons DM, Slater CJ, De Jesus V, Tang L, Singh R, Goldford JE, Martin S, Huang Q, Francis EA, Habsid A, Climie R, Tieu D, Wei J, Li R, Tong AHY, Aregger M, Chan KS, Han H, Wang X, Mero P, Brumell JH, Finelli A, Ailles L, Bader G, Smolen GA, Kingsbury GA, Hart T, Kung C, Moffat J. Functional genomic landscape of cancer-intrinsic evasion of killing by T cells. Nature. 2020;586(7827). doi: 10.1038/s41586-020-2746-2.

22. McGranahan N, Furness AJ, Rosenthal R, Ramskov S, Lyngaa R, Saini SK, Jamal-Hanjani M, Wilson GA, Birkbak NJ, Hiley CT, Watkins TB, Shafi S, Murugaesu N, Mitter R, Akarca AU, Linares J, Marafioti T, Henry JY, Van Allen EM, Miao D, Schilling B, Schadendorf D, Garraway LA, Makarov V, Rizvi NA, Snyder A, Hellmann MD, Merghoub T, Wolchok JD, Shukla SA, Wu CJ, Peggs KS, Chan TA, Hadrup SR, Quezada SA, Swanton C. Clonal neoantigens elicit T cell immunoreactivity and sensitivity to immune checkpoint blockade. Science. 2016;351(6280):1463–9. doi: 10.1126/science.aaf1490. PubMed PMID: 26940869.

23. Wolf Y, Bartok O, Patkar S, Eli GB, Cohen S, Litchfield K, Levy R, Jiménez-Sánchez A, Trabish S, Lee JS, Karathia H, Barnea E, Day CP, Cinnamon E, Stein I, Solomon A, Bitton L, Pérez-Guijarro E, Dubovik T, Shen-Orr SS, Miller ML, Merlino G, Levin Y, Pikarsky E, Eisenbach L, Admon A, Swanton C, Ruppin E, Samuels Y. UVB-Induced Tumor Heterogeneity Diminishes Immune Response in Melanoma. Cell. 2019;179(1):219–35.e21. doi: 10.1016/J.CELL.2019.08.032.

24. Melo M, Porter E, Zhang Y, Silva M, Li N, Dobosh B, Liguori A, Skog P, Landais E, Menis S, Sok D, Nemazee D, Schief WR, Weiss R, Irvine DJ. Immunogenicity of RNA Replicons Encoding HIV Env Immunogens Designed for Self-Assembly into Nanoparticles. Molecular therapy: the journal of the American Society of Gene Therapy. 2019;27(12):2080–90. Epub 20190819. doi: 10.1016/j.ymthe.2019.08.007. PubMed PMID: 31515132; PMCID: PMC6904793.

25. Li B, Luo X, Deng B, Wang J, McComb DW, Shi Y, Gaensler KM, Tan X, Dunn AL, Kerlin BA, Dong Y. An Orthogonal Array Optimization of Lipid-like Nanoparticles for mRNA Delivery in Vivo. Nano letters. 2015;15(12):8099–107. Epub 20151106. doi: 10.1021/acs.nanolett.5b03528. PubMed PMID: 26529392; PMCID: PMC4869688.

26. Yadav M, Jhunjhunwala S, Phung QT, Lupardus P, Tanguay J, Bumbaca S, Franci C, Cheung TK, Fritsche J, Weinschenk T, Modrusan Z, Mellman I, Lill JR, Delamarre L. Predicting immunogenic tumour mutations by combining mass spectrometry and exome sequencing. Nature 2014 515:7528. 2014;515(7528):572–6. doi: 10.1038/nature14001.

27. Castle JC, Kreiter S, Diekmann J, Lower M, van de Roemer N, de Graaf J, Selmi A, Diken M, Boegel S, Paret C, Koslowski M, Kuhn AN, Britten CM, Huber C, Tureci O, Sahin U. Exploiting the mutanome for tumor vaccination. Cancer research. 2012;72(5):1081–91. Epub 2012/01/13. doi: 0008-5472.CAN-11-3722[pii]10.1158/0008-5472.CAN-11-3722. PubMed PMID: 22237626.

28. Matsushita H, Vesely MD, Koboldt DC, Rickert CG, Uppaluri R, Magrini VJ, Arthur CD, White JM, Chen YS, Shea LK, Hundal J, Wendl MC, Demeter R, Wylie T, Allison JP, Smyth MJ, Old LJ, Mardis ER, Schreiber RD. Cancer exome analysis reveals a T-cell-dependent mechanism of cancer immunoediting. Nature. 2012;482(7385):400–4. doi: 10.1038/nature10755. PubMed PMID: 22318521; PMCID: 3874809.

29. DuPage M, Dooley AL, Jacks T. Conditional mouse lung cancer models using adenoviral or lentiviral delivery of Cre recombinase. Nature protocols. 2009;4(7):1064–72. doi: 10.1038/nprot.2009.95. PubMed PMID: 19561589; PMCID: 2757265.

30. Ben-David U, Siranosian B, Ha G, Tang H, Oren Y, Hinohara K, Strathdee CA, Dempster J, Lyons NJ, Burns R, Nag A, Kugener G, Cimini B, Tsvetkov P, Maruvka YE, O’Rourke R, Garrity A, Tubelli AA, Bandopadhayay P, Tsherniak A, Vazquez F, Wong B, Birger C, Ghandi M, Thorner AR, Bittker JA, Meyerson M, Getz G, Beroukhim R, Golub TR. Genetic and transcriptional evolution alters cancer cell line drug response. Nature 2018 560:7718. 2018;560(7718):325–30. doi: 10.1038/s41586-018-0409-3.

31. Luksza M, Riaz N, Makarov V, Balachandran VP, Hellmann MD, Solovyov A, Rizvi NA, Merghoub T, Levine AJ, Chan TA, Wolchok JD, Greenbaum BD. A neoantigen fitness model predicts tumor response to checkpoint blockade immunotherapy. Nature. 2017;551(7681):517. doi: 10.1038/NATURE24473.

32. Nielsen M, Lundegaard C, Worning P, Lauemøller SL, Lamberth K, Buus S, Brunak S, Lund O. Reliable prediction of T-cell epitopes using neural networks with novel sequence representations. Protein science: a publication of the Protein Society. 2003;12(5):1007–17. doi: 10.1110/PS.0239403.

33. Andreatta M, Nielsen M. Gapped sequence alignment using artificial neural networks: application to the MHC class I system. Bioinformatics. 2016;32(4):511–7. doi: 10.1093/bioinformatics/btv639. PubMed PMID: 26515819.

34. Hildner K, Edelson BT, Purtha WE, Diamond M, Matsushita H, Kohyama M, Calderon B, Schraml BU, Unanue ER, Diamond MS, Schreiber RD, Murphy TL, Murphy KM. Batf3 deficiency reveals a critical role for CD8alpha+ dendritic cells in cytotoxic T cell immunity. Science. 2008;322(5904):1097–100. doi: 10.1126/science.1164206. PubMed PMID: 19008445; PMCID: 2756611.

35. Spranger S, Bao R, Gajewski TF. Melanoma-intrinsic beta-catenin signalling prevents anti-tumour immunity. Nature. 2015;523(7559):231–5. doi: 10.1038/nature14404. PubMed PMID: 25970248.

36. Duong E, Fessenden TB, Lutz E, Dinter T, Yim L, Blatt S, Bhutkar A, Wittrup KD, Spranger S. Type I interferon activates MHC class I-dressed CD11b(+) conventional dendritic cells to promote protective anti-tumor CD8(+) T cell immunity. Immunity. 2021. doi: 10.1016/j.immuni.2021.10.020. PubMed PMID: 34800368.

37. Tussiwand R, Lee WL, Murphy TL, Mashayekhi M, Kc W, Albring JC, Satpathy AT, Rotondo JA, Edelson BT, Kretzer NM, Wu X, Weiss LA, Glasmacher E, Li P, Liao W, Behnke M, Lam SSK, Aurthur CT, Leonard WJ, Singh H, Stallings CL, David Sibley L, Schreiber RD, Murphy KM. Compensatory dendritic cell development mediated by BATF-IRF interactions. Nature. 2012;490(7421):502. doi: 10.1038/NATURE11531.

38. Bullock TNJ, Mullins DW, Engelhard VH. Antigen Density Presented By Dendritic Cells In Vivo Differentially Affects the Number and Avidity of Primary, Memory, and Recall CD8+ T Cells. The Journal of Immunology. 2003;170(4):1822–9. doi: 10.4049/JIMMUNOL.170.4.1822.

39. Westcott PMK, Sacks NJ, Schenkel JM, Ely ZA, Smith O, Hauck H, Jaeger AM, Zhang D, Backlund CM, Beytagh MC, Patten JJ, Elbashir R, Eng G, Irvine DJ, Yilmaz OH, Jacks T. Low neoantigen expression and poor T-cell priming underlie early immune escape in colorectal cancer. Nature Cancer 2021 2:10. 2021;2(10):1071–85. doi: 10.1038/s43018-021-00247-z.

40. Ferris ST, Wu R, Durai V, Murphy TL, Murphy KM. cDC1 prime and receive help from CD4 T cells to promote anti-tumor responses. The Journal of Immunology. 2020;204(1 Supplement).

41. Mailliard RB, Egawa S, Cai Q, Kalinska A, Bykovskaya SN, Lotze MT, Kapsenberg ML, Storkus WJ, Kalinski P. Complementary Dendritic Cell–activating Function of CD8+ and CD4+ T Cells: Helper Role of CD8+ T Cells in the Development of T Helper Type 1 Responses. The Journal of experimental medicine. 2002;195(4):473. doi: 10.1084/JEM.20011662.

42. Hernandez MGH, Shen L, Rock KL. CD40-CD40 Ligand Interaction between Dendritic Cells and CD8+ T Cells Is Needed to Stimulate Maximal T Cell Responses in the Absence of CD4+ T Cell Help. The Journal of Immunology. 2007;178(5):2844–52. doi: 10.4049/JIMMUNOL.178.5.2844.

43. Yi R, Chen E, Roberts EW, Krummel MF, Kathrin Serwas N. Impact of protein identity on tumor-associated antigen uptake into infiltrating immune cells: A comparison of different fluorescent proteins as model antigens. PloS one. 2022;17(8). doi: 10.1371/JOURNAL.PONE.0272857.

44. Carenza C, Calcaterra F, Oriolo F, Di Vito C, Ubezio M, Porta DMG, Mavilio D, Bella DS. Costimulatory molecules and immune checkpoints are differentially expressed on different subsets of dendritic cells. Frontiers in Immunology. 2019;10(JUN):1325. doi: 10.3389/FIMMU.2019.01325/BIBTEX.

45. Kotaka M, Ruedl C. Cutting Edge: Granulocyte-Macrophage Colony-Stimulating Factor Is the Major CD8(+) T Cell-Derived Licensing Factor for Dendritic Cell Activation. Article in The Journal of Immunology. 2010. doi: 10.4049/jimmunol.0903873.

46. Van Mierlo GJD, Boer DAT, Medema JP, Van Der Voort EIH, Fransen MF, Offringa R, Melief CJM, Toes REM. CD40 stimulation leads to effective therapy of CD40-tumors through induction of strong systemic cytotoxic T lymphocyte immunity. Proceedings of the National Academy of Sciences of the United States of America. 2002;99(8):5561. doi: 10.1073/PNAS.082107699.

47. Sandin LC, Orlova A, Gustafsson E, Ellmark P, Tolmachev V, Tötterman TH, Mangsbo SM. Locally delivered CD40 agonist antibody accumulates in secondary lymphoid organs and eradicates experimental disseminated bladder cancer. Cancer immunology research. 2014;2(1):80–90. doi: 10.1158/2326-6066.CIR-13-0067.

48. Morrison AH, Diamond MS, Hay CA, Byrne KT, Vonderheide RH. Sufficiency of CD40 activation and immune checkpoint blockade for T cell priming and tumor immunity. Proceedings of the National Academy of Sciences of the United States of America. 2020;117(14):8022–31. doi: 10.1073/PNAS.1918971117/SUPPL_FILE/PNAS.1918971117.SAPP.PDF.

49. Gejman RS, Chang AY, Jones HF, Dikun K, Hakimi AA, Schietinger A, Scheinberg DA. Rejection of immunogenic tumor clones is limited by clonal fraction. eLife. 2018;7. doi: 10.7554/ELIFE.41090.

50. Wu R, Murphy KM. DCs at the center of help: Origins and evolution of the three-cell-type hypothesis. The Journal of experimental medicine. 2022;219(7). Epub 20220511. doi: 10.1084/jem.20211519. PubMed PMID: 35543702; PMCID: PMC9098650.

51. Wu R, Murphy KM. DCs at the center of help: Origins and evolution of the threecell-type hypothesis. The Journal of experimental medicine. 2022;219(7). doi: 10.1084/JEM.20211519/213211.

52. Ferris ST, Durai V, Wu R, Theisen DJ, Ward JP, Bern MD, Davidson JT, Bagadia P, Liu T, Briseño CG, Li L, Gillanders WE, Wu GF, Yokoyama WM, Murphy TL, Schreiber RD, Murphy KM. cDC1 prime and are licensed by CD4+ T cells to induce anti-tumour immunity. Nature. 2020;584(7822). doi: 10.1038/s41586-020-2611-3.

53. Gubin MM, Zhang X, Schuster H, Caron E, Ward JP, Noguchi T, Ivanova Y, Hundal J, Arthur CD, Krebber WJ, Mulder GE, Toebes M, Vesely MD, Lam SS, Korman AJ, Allison JP, Freeman GJ, Sharpe AH, Pearce EL, Schumacher TN, Aebersold R, Rammensee HG, Melief CJ, Mardis ER, Gillanders WE, Artyomov MN, Schreiber RD. Checkpoint blockade cancer immunotherapy targets tumour-specific mutant antigens. Nature. 2014;515(7528):577–81. doi: 10.1038/nature13988. PubMed PMID: 25428507; PMCID: PMC4279952.

54. Burger ML, Cruz AM, Crossland GE, Gaglia G, Ritch CC, Blatt SE, Bhutkar A, Canner D, Kienka T, Tavana SZ, Barandiaran AL, Garmilla A, Schenkel JM, Hillman M, de los Rios Kobara I, Li A, Jaeger AM, Hwang WL, Westcott PMK, Manos MP, Holovatska MM, Hodi FS, Regev A, Santagata S, Jacks T. Antigen dominance hierarchies shape TCF1+ progenitor CD8 T cell phenotypes in tumors. Cell. 2021;184(19):4996–5014.e26. doi: 10.1016/J.CELL.2021.08.020.

55. Kotturi MF, Scott I, Wolfe T, Peters B, Sidney J, Cheroutre H, von Herrath MG, Buchmeier MJ, Grey H, Sette A. Naive Precursor Frequencies and MHC Binding Rather Than the Degree of Epitope Diversity Shape CD8 + T Cell Immunodominance. The Journal of Immunology. 2008;181(3). doi: 10.4049/jimmunol.181.3.2124.

56. Willimsky G, Blankenstein T. Sporadic immunogenic tumours avoid destruction by inducing T-cell tolerance. Nature 2005 437:7055. 2005;437(7055):141–6. doi: 10.1038/nature03954.

57. Schietinger A, Philip M, Krisnawan VE, Chiu EY, Delrow JJ, Basom RS, Lauer P, Brockstedt DG, Knoblaugh SE, Hämmerling GJ, Schell TD, Garbi N, Greenberg PD. Tumor-Specific T Cell Dysfunction Is a Dynamic Antigen-Driven Differentiation Program Initiated Early during Tumorigenesis. Immunity. 2016;45(2):389–401. doi: 10.1016/J.IMMUNI.2016.07.011/ATTACHMENT/39522132-2F2F-4C0D-A65A-EF8F2F1C6480/MMC5.XLSX.

58. Spiotto MT, Yu P, Rowley DA, Nishimura MI, Meredith SC, Gajewski TF, Fu YX, Schreiber H. Increasing tumor antigen expression overcomes “ignorance” to solid tumors via crosspresentation by bone marrow-derived stromal cells. Immunity. 2002;17(6):737–47. PubMed PMID: 12479820.

59. Frentsch M, Stark R, Matzmohr N, Meier S, Durlanik S, Schulz AR, Stervbo U, Jürchott K, Gebhardt F, Heine G, Reuter MA, Betts MR, Busch D, Thiel A. CD40L expression permits CD8+ T cells to execute immunologic helper functions. Blood. 2013;122(3):405. doi: 10.1182/BLOOD-2013-02-483586.

60. Wong KL, Lew FC, MacAry PA, Kemeny DM. CD40L-expressing CD8 T cells prime CD8alpha(+) DC for IL-12p70 production. European journal of immunology. 2008;38(8):2251–62. doi: 10.1002/EJI.200838199.

61. Pfirschke C, Engblom C, Rickelt S, Cortez-Retamozo V, Garris C, Pucci F, Yamazaki T, Poirier-Colame V, Newton A, Redouane Y, Lin YJ, Wojtkiewicz G, Iwamoto Y, Mino-Kenudson M, Huynh TG, Hynes RO, Freeman GJ, Kroemer G, Zitvogel L, Weissleder R, Pittet MJ. Immunogenic Chemotherapy Sensitizes Tumors to Checkpoint Blockade Therapy. Immunity. 2016;44(2):343–54. doi: 10.1016/j.immuni.2015.11.024. PubMed PMID: 26872698; PMCID: PMC4758865.

62. Starbeck-Miller GR, Xue HH, Harty JT. IL-12 and type I interferon prolong the division of activated CD8 T cells by maintaining high-affinity IL-2 signaling in vivo. The Journal of experimental medicine. 2014;211(1):105. doi: 10.1084/JEM.20130901.

63. Horton BL, Morgan DM, Momin N, Zagorulya M, Torres-Mejia E, Bhandarkar V, Wittrup KD, Love JC, Spranger S. Lack of CD8(+) T cell effector differentiation during priming mediates checkpoint blockade resistance in non-small cell lung cancer. Sci Immunol. 2021;6(64):eabi8800. doi: 10.1126/sciimmunol.abi8800. PubMed PMID: 34714687.

64. Momin N, Mehta NK, Bennett NR, Ma L, Palmeri JR, Chinn MM, Lutz EA, Kang B, Irvine DJ, Spranger S, Wittrup KD. Anchoring of intratumorally administered cytokines to collagen safely potentiates systemic cancer immunotherapy. Science translational medicine. 2019;11(498). doi: 10.1126/scitranslmed.aaw2614. PubMed PMID: 31243150.

65. Castellino F, Huang AY, Altan-Bonnet G, Stoll S, Scheinecker C, Germain RN. Chemokines enhance immunity by guiding naive CD8+ T cells to sites of CD4+ T cell-dendritic cell interaction. Nature. 2006;440(7086):890–5. doi: 10.1038/NATURE04651.

66. Spranger S, Dai D, Horton B, Gajewski TF. Tumor-Residing Batf3 Dendritic Cells Are Required for Effector T Cell Trafficking and Adoptive T Cell Therapy. Cancer cell. 2017;31(5):711–23 e4. doi: 10.1016/j.ccell.2017.04.003. PubMed PMID: 28486109.

67. Gardner A, de Mingo Pulido A, Hanggi K, Bazargan S, Onimus A, Kasprzak A, Conejo-Garcia JR, Rejniak KA, Ruffell B. TIM-3 blockade enhances IL-12-dependent antitumor immunity by promoting CD8(+) T cell and XCR1(+) dendritic cell spatial colocalization. Journal for immunotherapy of cancer. 2022;10(1). doi: 10.1136/jitc-2021-003571. PubMed PMID: 34987021; PMCID: PMC8734033.

68. Wroblewska A, Dhainaut M, Ben-Zvi B, Rose SA, Park ES, Amir ED, Bektesevic A, Baccarini A, Merad M, Rahman AH, Brown BD. Protein Barcodes Enable High-Dimensional Single-Cell CRISPR Screens. Cell. 2018;175(4):1141–55 e16. Epub 20181018. doi: 10.1016/j.cell.2018.09.022. PubMed PMID: 30343902; PMCID: PMC6319269.

69. Angelova M, Mlecnik B, Vasaturo A, Bindea G, Fredriksen T, Lafontaine L, Buttard B, Morgand E, Bruni D, Jouret-Mourin A, Hubert C, Kartheuser A, Humblet Y, Ceccarelli M, Syed N, Marincola FM, Bedognetti D, Van den Eynde M, Galon J. Evolution of Metastases in Space and Time under Immune Selection. Cell. 2018;175(3):751–65 e16. doi: 10.1016/j.cell.2018.09.018. PubMed PMID: 30318143.

70. Alcantara-Hernandez M, Leylek R, Wagar LE, Engleman EG, Keler T, Marinkovich MP, Davis MM, Nolan GP, Idoyaga J. High-Dimensional Phenotypic Mapping of Human Dendritic Cells Reveals Interindividual Variation and Tissue Specialization. Immunity. 2017;47(6):1037–50 e6. doi: 10.1016/j.immuni.2017.11.001. PubMed PMID: 29221729; PMCID: PMC5738280.

